# Innate immune sensing via the cGAS-STING pathway restricts extrachromosomal DNA–driven tumorigenesis

**DOI:** 10.64898/2025.12.31.697191

**Authors:** Tuo Li, Qing-Lin Yang, Kailiang Qiao, Anli Zhang, Chenglong Sun, Huocong Huang, Paul S. Mischel, Sihan Wu, Zhijian J. Chen

**Author notes:** These authors contributed equally to this work (Tuo Li, Qing-Lin Yang).

## Abstract

Extrachromosomal DNAs (ecDNAs) are circular DNA fragments frequently found in human cancers, where they amplify oncogenes, drive tumor heterogeneity, and promote therapy resistance and poor prognosis. Despite their prevalence, how ecDNAs interact with the immune system remains poorly understood. Here, we show that the cytosolic DNA sensor cGAS detects ecDNA fragments in the cytoplasm and activates the innate immune response. cGAS and STING are frequently silenced in ecDNA^+^ tumors through promoter hypermethylation. Restoring cGAS or STING in human and murine ecDNA^+^ cancer cells reactivates innate immune signaling and selectively suppresses ecDNA^+^ tumor growth in an immunocompetent mouse model. Using two ecDNA biogenesis models, we show that the cGAS-STING pathway restricts *de novo* ecDNA formation. Together, our findings identify innate immune sensing as a natural barrier to ecDNA-driven oncogenesis and establish cGAS-STING reactivation as a therapeutic strategy for ecDNA^+^ cancers.

**Highlights:**

- The cGAS-STING pathway is frequently silenced in ecDNA^+^ tumors
- Restoration of cGAS in ecDNA^+^ cells activates innate immune responses
- cGAS expression suppresses ecDNA^+^ tumor growth *in vivo*
- The cGAS-STING pathway restricts *de novo* ecDNA biogenesis

## Introduction

Extrachromosomal DNAs (ecDNAs) are large, acentric circular chromatin fragments frequently observed in cancer cells^1,2^. These structures, typically spanning hundreds of kilobases, arise from focal genomic rearrangements and exist outside the linear chromosomes. ecDNAs often carry amplified oncogenes such as *MYC*, *EGFR*, or *KRAS*^3^, as well as immunomodulatory genes and regulatory elements^4^ driving overexpression of tumor promoting genes through increased copy number, highly accessible chromatin, and elevated transcriptional activity^5^. They can also harbor enhancer elements that interact *in trans* with other ecDNAs^6^ or chromosomal loci^7^ to promote oncogenic transcriptional networks. Because ecDNAs segregate randomly during mitosis^8^ and can co-segregate with other ecDNA species^9^, they create extensive copy number heterogeneity within tumors, fueling rapid clonal evolution and drug resistance^3,8,10–12^. Indeed, ecDNA abundance strongly correlates with tumor aggressiveness and poor clinical outcomes across diverse malignancies^13^, underscoring the urgent need for strategies that specifically target ecDNA-positive (ecDNA⁺) cancers.

Despite extensive characterization of the genomic and transcriptional properties of ecDNAs, how these structures interact with immune defense mechanisms remains unknown. ecDNAs contain more accessible double-stranded DNA (dsDNA) that differs topologically from chromosomal DNA^5^, yet cancer cells carrying large amounts of ecDNA rarely display hallmarks of innate immune activation^14,15^. This raises a fundamental question: are ecDNAs invisible to immune surveillance, or have tumors evolved strategies to silence or evade their detection? Addressing this question could reveal a missing link between ecDNA and tumor immune evasion^16^.

The cytosolic DNA sensor *c*yclic *G*MP–*A*MP *s*ynthase (cGAS) provides an appealing candidate for immune surveillance of tumors. cGAS detects dsDNA in the cytoplasm independent of sequence^17^ and catalyzes the formation of a cyclic dinucleotide—2’3’-cyclic GMP-AMP (cGAMP)^18,19^. cGAMP is a second messenger that binds to the adaptor protein *ST*imulator of *IN*terferon *G*enes (STING), which resides on the endoplasmic reticulum membrane and translocates to the Golgi apparatus upon activation^20^. STING recruits the kinases TBK1 and IKK, leading to phosphorylation of IRF3 and NF-κB and subsequent induction of type I interferons and inflammatory cytokines. This pathway provides a crucial first line of defense against viral and self-DNA in the cytoplasm^21^, and dysregulation of cGAS–STING signaling can shape tumor immunogenicity and therapeutic response^22^.

Emerging transcriptomic analyses suggest that immune signaling components are frequently downregulated in ecDNA⁺ tumors^14,15^. Yet the mechanism and consequence of this suppression remain unclear. In this study, we define how the cGAS–STING pathway interacts with ecDNA in cancer. We analyze its expression landscape across large patient cohorts, uncover its frequent silencing via promoter hypermethylation, and demonstrate that restoring cGAS–STING signaling revives innate immunity and selectively inhibits ecDNA⁺ tumor growth. Finally, using cellular models of ecDNA formation, we reveal that active cGAS–STING signaling constrains *de novo* ecDNA biogenesis. Together, our findings establish innate immune sensing by the cGAS-STING pathway as a natural barrier to ecDNA-driven oncogenesis.

## Results

### The cGAS-STING pathway is frequently defective in ecDNA^+^ cancer cells

Analysis of bulk mRNA sequencing data from overlapping TCGA and PCAWG (AmpliconRepo^23^) cohorts revealed that *cGAS* (MB21D1) expression is markedly lower than that of *STING* (TMEM173), *RIG*-I (DDX58), *MDA5* (IFIH1), *MAVS*, *TBK1*, and other nucleic acid–sensing pathway genes (p < 0.0001, Figure S1A). The overall innate immune landscape appeared comparable among tumors with ecDNA (n = 189), with other focal amplifications (n = 117), and with no detected focal amplifications (n = 315). To confirm this finding at the protein level, we examined cGAS and STING expression levels by immunoblots in a panel of patient-derived ecDNA^+^ cancer cells (Figure 1A). cGAS was not detected in the colorectal adenocarcinoma COLO320 isogenic cell lines—DM (ecDNA^+^) and HSR (ecDNA^-^), the glioblastoma GBM39 isogenic cell lines—KT (ecDNA^+^) and HSR (ecDNA^-^), or neuroblastoma ecDNA^+^ cell lines CHP212 and Kelly. The gastric carcinoma SNU16 cell line expressed cGAS but not STING. The prostate adenocarcinoma PC3 cell line expressed both cGAS and STING. When we transfected these cells with herring testis derived DNA (HT-DNA), they all showed blunted transcriptional activation of *Interferon β* (*IFNβ*) or *CXCL10*, compared to THP1 cells (p < 0.0001), which are known to express these cytokines (Figure 1B). Although PC3 cells express both cGAS and STING, *IFNβ* and *CXCL10* were only modestly induced in them in response to HT-DNA compared to in THP1 cells (p < 0.0001), suggesting additional defects in the signaling pathway in this cell line. RNA sequencing of these cell lines revealed that the *cGAS*^+^ and *cGAS*^-^ groups can be separated by a threshold of TPM of 1 (Figure S1B). Applying this threshold, we found that in 22% of TCGA tumors, *cGAS* expression can be deemed negative (Figure S1A). In ecDNA^+^ tumors, the *cGAS*^-^percentage is similar—19% (Figure S1A). Because tumor samples contain *cGAS*-expressing non-malignant cells such as fibroblasts and macrophages, ∼19% may underestimate the portion of cGAS-low tumors. While basal levels of RIG-I and MDA5 are typically low prior to stimulation^24^, pan-cancer analysis indicates that *cGAS* levels in tumors are even lower. Therefore, these analyses showed that the cytosolic DNA sensing pathway is defective in a significant portion of human tumors, including those with ecDNA. To understand why *cGAS* or *STING* expression was lost in these cancer cells, we measured the methylation status of their genomic loci by bisulfite DNA sequencing. We first identified a CpG island that covers the promoter and the first exon of *cGAS* locus and found that it was hypermethylated in the COLO320, GBM, and neuroblastoma cell lines, correlating with the absence of cGAS in these cells (Figure 1C). These results show that cGAS expression is regulated by promoter DNA methylation, consistent with other studies^25,26^. In contrast, CpG methylation at the *STING* promoter locus does not always correlate with STING protein levels (Figure S1C), suggesting other levels of regulation. Therefore, epigenetic silencing may underlie the loss of *cGAS* expression in cancer cells.

**Figure 1.**
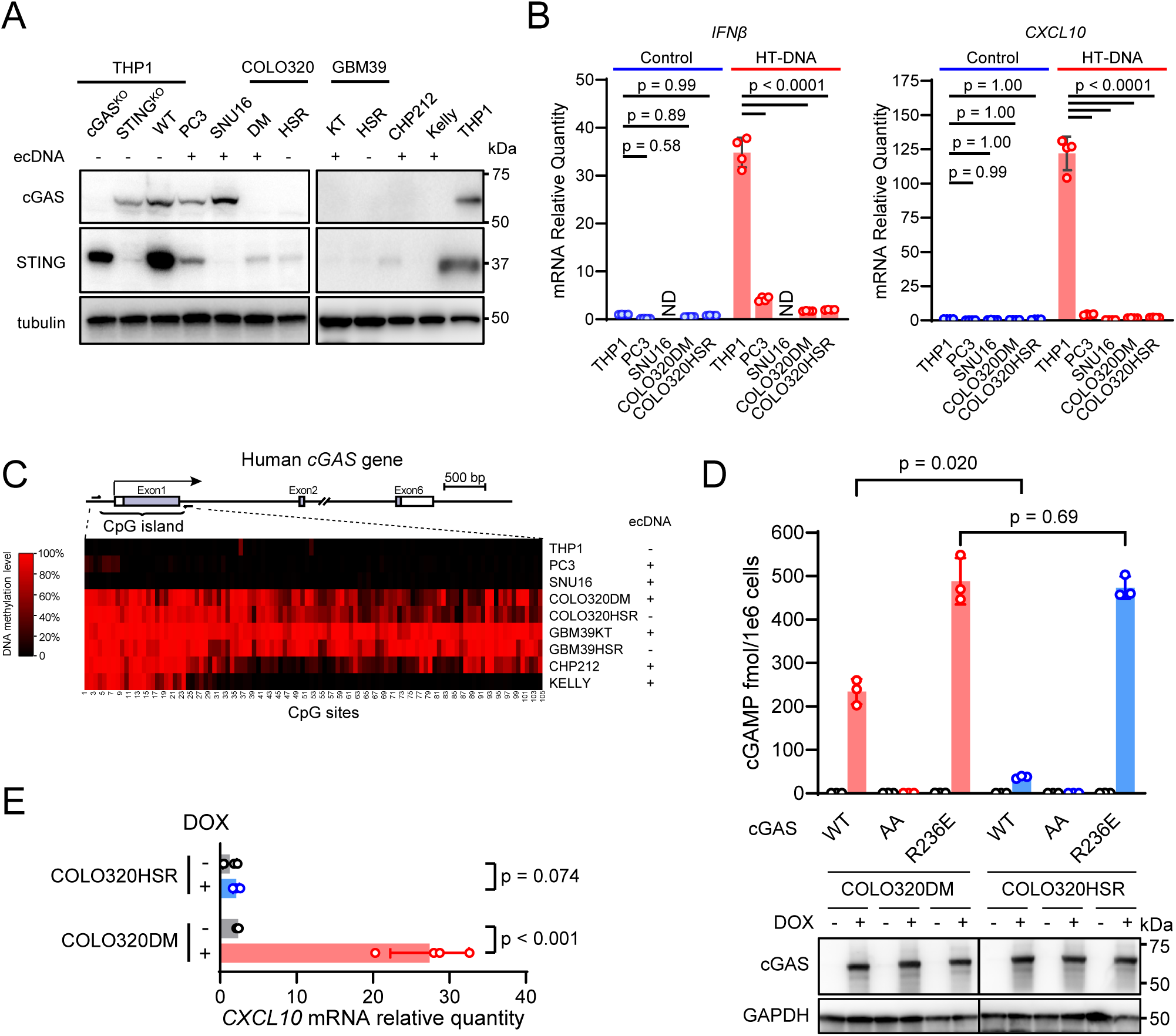
Immune activation by restoring the cGAS-STING pathway in human ecDNA^+^ cancer cells. (**A**) Immunoblotting images show the protein levels of cGAS and STING in a panel of ecDNA^+^ cancer cells with their ecDNA status indicated. Positive control: THP1; negative controls: THP1 ΔcGAS and THP1 ΔSTING cells. (**B**) The cancer cell lines were transfected with 1 µg/mL HT-DNA for 6 hours and the mRNA levels of *IFNβ* and *CXCL10* were quantified by RT-qPCR. ND: not detected. (**C**) The schematics show the location of the CpG island in Exon 1 of human *cGAS* gene. The heatmaps show the DNA methylation levels of individual CpG sites as measured by bisulfite sequencing. (**D**) COLO320DM (ecDNA^+^) and COLO320HSR (ecDNA^-^) cells were stably transduced with lentiviruses harboring cGAS expression constructs controlled by a tetracycline-inducible promoter. Wild-type (WT), the catalytically dead mutant G213A/S214A (AA), and the constitutively de-repressed mutant (R236E) of cGAS are indicated. After induction with 0.1 µg/mL doxycycline for 24 hours, cGAMP levels were measured by mass spectrometry (*upper*) and cGAS protein levels determined by immunoblotting (*bottom*). (**E**) *CXCL10* mRNA levels were measured by RT-qPCR in COLO320DM and COLO320HSR cells expressing wild-type cGAS.

### Restoration of cGAS in human ecDNA^+^ cancer cells activates innate immune response

Given that the cGAS-STING pathway is frequently defective in ecDNA^+^ cancer cells, we tested if the immune response could be restored by expressing cGAS. We generated stable SNU16 cell lines that express wild-type STING or the cGAMP-binding-deficient STING mutant, R238A/Y240A, under the tetracycline-inducible promoter. After doxycycline induction, there was an eight-fold increase in *CXCL10* mRNA levels in cells expressing wild-type STING (p < 0.001), but no elevation in cells expressing the mutant STING (Figure S2A&B, p = 0.93). As a positive control, the STING agonist MSA2 strongly induced *CXCL10* transcription in SNU16 cells with wild-type STING. The data shows that transient restoration of STING expression in the ecDNA^+^ SNU16 cells leads to cGAS-STING pathway activation.

Similarly, we used the tetracycline system to restore cGAS expression in the COLO320 isogenic pair. In COLO320DM cells that express wild-type cGAS, we detected 234 fmol cGAMP per million cells, while in COLO320HSR cells, cGAS induced 37 fmol cGAMP (Figure 1D, p = 0.020). As a positive control, the histone-binding-deficient mutant R236E of cGAS, which is activated by nuclear genomic DNA^27^, produced strong and equivalent levels of cGAMP in both COLO320DM and COLO320HSR cell lines (p = 0.69). As a negative control, the catalytically dead mutant (G213A/S214A; denoted as AA) of cGAS did not produce cGAMP in either cell line (Figure 1D). Doxycycline-induced expression of wild-type cGAS in ecDNA^-^ human cell lines, BJ-5ta or THP1, did not generate any detectable cGAMP, indicating that overexpression alone does not activate wild-type cGAS, presumably because these cells lack cytosolic DNA (Figure S2C). Furthermore, we tested the inducible expression of cGAS in glioblastoma cell lines GBM39KT (ecDNA^+^) and GBM39HSR (ecDNA^-^). Transient expression of wild-type cGAS produced cGAMP only in GBM39KT cells but not in GBM39HSR cells (Figure S2D). In COLO320DM cells but not in COLO320HSR cells, doxycycline also induced the expression of *CXCL10* (Figure 1E). Therefore, cGAS can be activated by endogenous DNA specifically in human ecDNA^+^ cancer cells.

### cGAS is activated by cytosolic ecDNA

Previous studies show that ecDNA clustered inside the nucleus of interphase cells in structures deemed ecDNA hubs^6^. During cell division, ecDNAs also cluster and piggyback on chromosomes by a less clear mechanism and segregate into daughter cells in a random manner^8^. ecDNA is rarely found in the cytoplasm by fluorescent *in situ* hybridization (FISH). We extracted DNA from the cytosolic fraction of COLO320DM and COLO320HSR cells and measured by quantitative PCR (qPCR) levels of DNA sequences representing ecDNA (*MYC1* and *MYC2*), nuclear genome (*B2M*), or mitochondria (*Dloop* and *ND1*, Figure S3A). The result revealed higher ecDNA levels in the cytosol of COLO320DM cells compared to those detected in the cytosol of COLO320HSR cells (p = 0.012 and, 0.015 for *MYC1* and *MYC2*, respectively), while the levels of nuclear genomic DNA or mitochondria DNA in the cytosol were not significantly different (p = 0.52, 0.36 and 0.054 for *B2M*, *Dloop* and *ND1*, respectively) (Figure S3B). Although the total ecDNA levels are also higher in the COLO320DM than in the COLO320HSR cells (1.86x and 1.82x for *MYC1* and *MYC2*, respectively), the fold enrichment is even higher in the cytosol (2.74x and 2.49x for *MYC1* and *MYC2*, respectively), suggesting selective enrichment of ecDNA sequence in the cytosol of COLO320DM cells. Therefore, low levels of ecDNA are present in the cytosol and can activate cGAS. A recent study reported that acentric chromosome fragments cluster upon mitotic entry via the CIP2A-TOPBP1 complex^28^. We tested whether CIP2A may also play a role in restraining ecDNA from the cytosol after mitosis, thus preventing cGAS activation. COLO320DM cells deficient in CIP2A indeed showed increased cGAS activation compared to the control cells when wild-type cGAS was expressed (Figure S3C, p = 0.011). As a control, the cGAS^R236E^ mutant was not affected by CIP2A knockdown (p = 0.56). In COLO320HSR cells, CIP2A knockdown also moderately increased cGAS activity (p = 0.001). Therefore, loss of CIP2A could promote cGAS activation by ecDNA.

To investigate how cGAS is activated in ecDNA^+^ cells, we visualized cGAS via a C-terminal GFP tag. In ∼10% COLO320DM cells, cGAS-GFP shows numerous foci in the cytosol (Figure 2A). In contrast, cGAS-GFP is diffused in the cytoplasm and nucleus of COLO320HSR cells. cGAS-GFP foci in COLO320DM cells recovered rapidly after photobleaching, indicating that they were liquid-phase separated condensates, a hallmark of cGAS activation^29^ (Figure 2B). Combining FISH and immunostaining, we could detect *MYC* sequence overlapping with cytosolic cGAS-GFP puncta, supporting that cGAS was activated by ecDNA in the cytoplasm (Figure 1C). To further confirm that cGAS is activated in the cytoplasmic puncta, we implemented the tripartite GFP complementation method previously established to detect cGAS activation (Figure 2D)^30^. cGAS was tagged with two small fragments of the enhanced monomeric GFP—G_10_ and G_11_ and co-expressed under the tetracycline promoter with the remainder of GFP—GFP-OPT_1-9_. After tetracycline induction, we observed GFP puncta in the cytosol, indicating cGAS oligomerization, but not in the nucleus of COLO320DM cells (Figure 2D). Little GFP signal was observed in COLO320HSR cells. These results suggest that cGAS is activated in the cytoplasmic puncta of COLO320DM cells. We also measured cGAS activity in subcellular fractions of COLO320DM cells by an *in vitro* assay following differential centrifugation. We observed higher cGAMP production in the cytoplasmic fractions of COLO320DM—S1 & S10, than in the nuclear fractions—P1 and P10 (Figure 2E). cGAS activity is low in all fractions of COLO320HSR cells. This result is consistent with the model that nuclear cGAS is tethered to and inhibited by nucleosomes^27,31–33^, and pinpoints to the cytoplasm where cGAS is activated in ecDNA^+^ cells.

**Figure 2.**
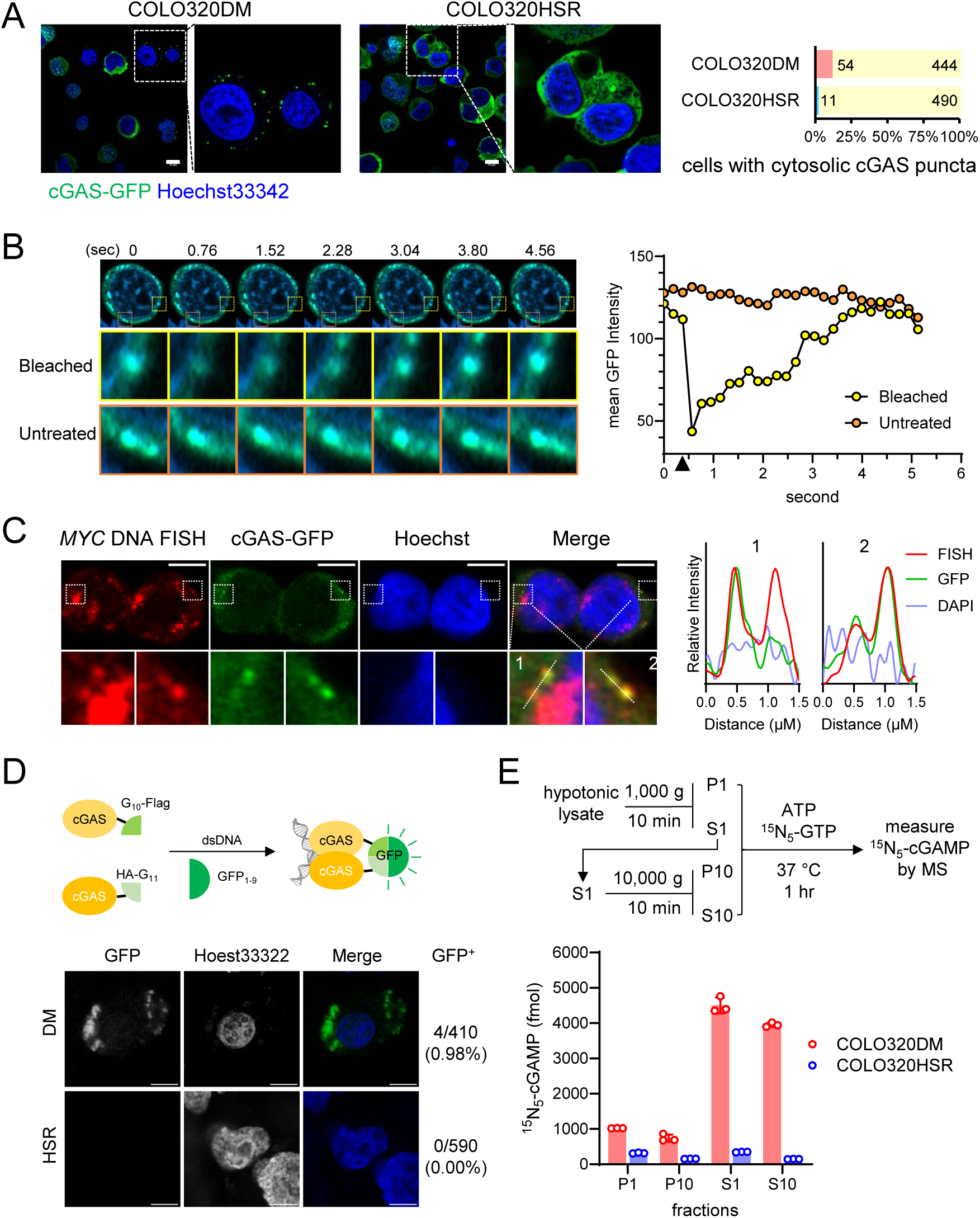
cGAS is activated by ecDNA in the cytoplasm of ecDNA^+^ cancer cells. (**A**) *(left)* Airyscan images of COLO320DM and COLO320HSR cells expressing cGAS-GFP (green) under the tetracycline-inducible promoter after doxycycline induction. DNA was stained by Hoechst 33342 (blue). Bars = 10 µm. *(right)* percentage of cells observed to have cytosolic cGAS puncta. (**B**) *(left)* Serial airyscan images show the recovery after targeted photobleaching of a cGAS-GFP punctum in the cytoplasm of COLO320DM cells (in yellow boxes). Orange boxes mark an untreated punctum for comparison. *(right)* mean GFP intensities of the selected cGAS-GFP puncta over time (photobleaching is marked by the black triangle). (**C**) *(left)* in COLO320DM cells, cGAS-GFP was immunostained with anti-GFP antibody and *MYC* ecDNA was visualized by DNA FISH. Bars = 5 µm. *(right)* relative intensities of FISH, GFP and DAPI along indicated dotted lines in the zoomed images. (**D**) A split GFP complementation assay for cGAS activation in COLO320DM and COLO320HSR cells. (*upper*) A schematic of the fluorescence reporter system; (*bottom*) Confocal microscopy images show GFP signals. Numbers indicate the percentage of GFP^+^ cells observed. Bars = 10 µm. (**E**) cGAS activity in COLO320DM or COLO320HSR cell lysates. (*upper*) The workflow of fractionation and the *in vitro* cGAS activity assay; (*bottom*) levels of ^15^N_5_-cGAMP as the product of cGAS activity in cell lysate fractions. P1 and S1, pellets and supernatants after centrifugation at 1000 x g, respectively. P10 and S10: centrifugation at 10,000 x g.

### Restoring cGAS expression activates innate immune response in mouse ecDNA^+^ cancer cells

Current studies of ecDNA are limited to cell models of human origin, which hinders the study of immune response to ecDNA^+^ tumors. To establish an immune competent mouse model of ecDNA^+^ tumors, we screened a library of single-cell clones from the *KPfC* (*Kras^LSL−G12D/+^Trp53^fl/fl^Pdx1^Cre/+^*) tumor, a genetically engineered mouse model (GEMM) of pancreatic ductal adenocarcinoma. We identified CT1BA5^34^ , a PDAC cell clone that spontaneously harbored ecDNA (Figure 3A and Figure S4A). Two subclonal cell lines were further purified from the CT1BA5 cell line: one cell line carries ecDNA that is *Kras* positive—CT1BA5-EC, and the other cell line carries the *Kras* gene amplification as homogenous stain region—CT1BA5-HSR. *Fluorescence in situ hybridization* (FISH) in metaphase confirmed the focal amplification status in these cell lines and sequencing showed that the *Kras* allele carries the G12D mutation (co-submitted manuscript by Qiao et al^35^). Other than the form of focal amplification, the two cell lines are similar in morphology and growth rate in cell culture. Both CT1BA5-EC and CT1BA5-HSR cells express *STING* but lack *cGAS* expression, and *cGAS* silencing correlated with CpG island hypermethylation in its promoter (Figure 3B and Figure S4B).

**Figure 3.**
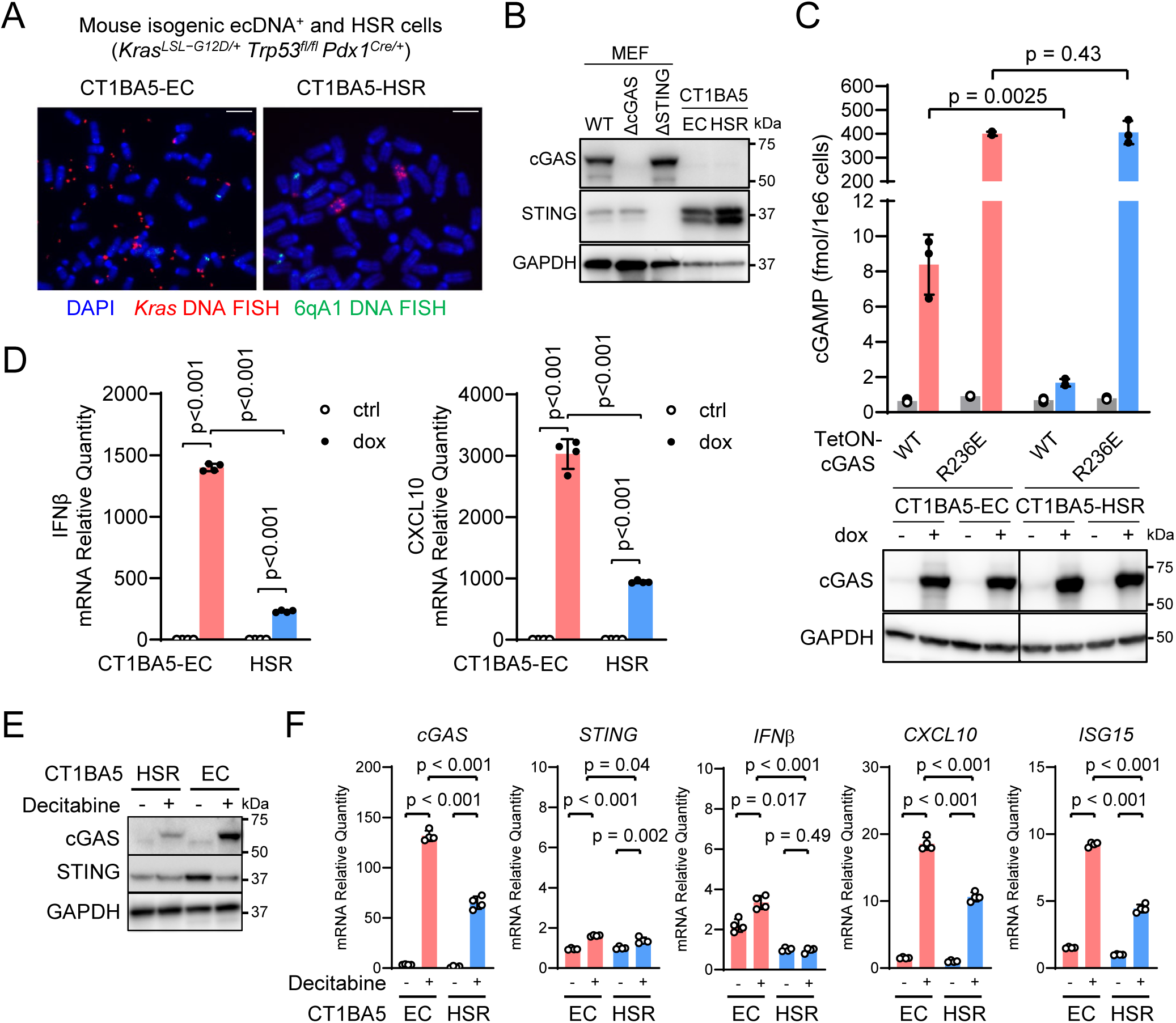
Immune activation by restoring the cGAS-STING pathway in murine ecDNA^+^ cancer cells. (**A**) Representative photos of metaphase spreads of CT1BA5-EC (ecDNA^+^) and CT1BA5-HSR (ecDNA^-^) cells. Bars = 5 µm. (**B**) Immunoblotting images show the protein levels of cGAS and STING in CT1BA5-EC and CT1BA5-HSR cells, with MEF cells serving as the positive control, and MEF ΔcGAS and MEF ΔSTING cells as negative controls. (**C**) CT1BA5-EC and CT1BA5-HSR cells were stably transduced with tetracycline-inducible cGAS lentiviral vectors: wild-type (WT), and the constitutively de-repressed mutant R236E. After induction with 0.1 µg/mL doxycycline for 24 hours, cGAMP levels were measured by mass spectrometry (upper) and cGAS protein levels by immunoblotting (bottom). (**D**) Quantify mRNA levels of *interferon β* and *cxcl10* by RT-qPCR in CT1BA5-EC and CT1BA5-HSR cells expressing wildtype cGAS. (**E&F**) CT1BA5EC and HSR cells were treated with 10 µM decitabine for 48 hrs. (E) Immunoblots show protein levels of cGAS and STING with or without treatment. (F) qPCR results show mRNA levels of *cGAS*, *STING*, *CXCL10*, *IFNβ*, and *ISG15*.

To test if cGAS can be activated in ecDNA^+^ murine tumor cells, we transiently induced cGAS expression with doxycycline. Like in ecDNA^+^ human tumor cells, ectopic expression of cGAS^WT^ in CT1BA5-EC cells produced cGAMP, although to a level lower than the histone-binding-deficient mutant cGAS^R236E^, but substantially higher than the cGAMP level in CT1BA5-HSR cells that express similar levels of WT cGAS (Figure 3C, p = 0.0025). Because CT1BA5-EC cells express *STING*, cGAS activation led to transcriptional activation of cytokines *CXCL10* and *IFNβ* (Figure 3D). Therefore, ectopically expressed cGAS can be activated in ecDNA^+^ mouse cancer cells. To test if endogenous cGAS can be re-activated by inhibiting DNA methylation, we treated CT1BA5 cell lines with a cytidine analog—decitabine, which inhibits DNA methyltransferases^36^. Decitabine treatment reduced DNA methylation of the CpG island of *cGAS* in both CT1BA5 cell lines and induced the expression of *cGAS* (Figure 3E&F, Figure S4C). Concomitantly, we observed transcriptional activation of *CXCL10*, *IFNβ*, and *ISG15* (Figure 3F), substantially more in the CT1BA5-EC cells than in the CT1BA5-HSR cells (p < 0.001), consistent with the outcome of the doxycycline induction experiment. Therefore, we can restore the functional cGAS-STING pathway and ecDNA sensing with the DNA methylation inhibitor decitabine.

### Ectopic cGAS selectively impedes ecDNA^+^ tumor growth

Both CT1BA5-EC and CT1BA5-HSR cells grew into tumors after grafting to immunodeficient (NSG) mice or immunocompetent C57BL/6 mice (Figure S5A). This is because the original CT1BA5 cell line was from a GEMM on the C57BL/6 background^34^. This unique property makes these cells amenable to immunology investigations. Although CT1BA5-EC and CT1BA5-HSR cells grow at the same rate *in vitro* ^35^, CT1BA5-EC tumors grow significantly faster than CT1BA5-HSR tumors (p = 0.033), and both are faster than the BMFA3 cell line from the same *KPfC* GEMM, in which *Kras* alleles are not amplified. Such growth differences highlight the role of *Kras* amplification in promoting tumorigenesis and suggest that copy number heterogeneity further fuels malignancy. We found that both EC and HSR tumors had low levels of immune cell infiltration^35^, and both were refractory to immune checkpoint blockades with the anti-PD-L1 antibody or with the anti-CTLA4 antibody (Figure S5B). CT1BA5-EC tumors were also resistant, and HSR tumors were partially resistant to the targeted therapy by the Kras^G12D^ inhibitor MRTX1133^37,38^ (Figure S5C), likely because of their high *Kras* allele copy numbers. In contrast, BMFA3 tumors are sensitive to MRTX1133. Notably, cGAMP can effectively inhibit the growth of both CT1BA5-EC and -HSR tumors (Figure S5D), indicating that activating the cGAS-STING pathway can induce a potent antitumor effect on these immunologically cold tumors.

Because cGAS can be selectively activated in ecDNA^+^ cells, we reason that ectopic cGAS expression would be a strategy to confer selectivity on ecDNA^+^ tumors over ecDNA^-^tumors. To test this hypothesis, we generated stable CT1BA5-EC and CT1BA5-HSR cells that express cGAS under a tetracycline-inducible promoter, and subcutaneously allografted C57BL/6 mice. The tumor-bearing mice were treated with doxycycline in drinking water to induce cGAS expression in the tumor cells. Remarkably, the growth of CT1BA5-EC tumors induced to express cGAS^WT^ was substantially inhibited compared to tumors without induction (Figure 4A&B, p < 0.0001). CT1BA5-EC tumors expressing the inactive cGAS^AA^ mutant were not affected by doxycycline induction (p = 0.65). Mice bearing CT1BA5-EC tumors that expressed cGAS^WT^ survived better than those that did not express cGAS or expressed cGAS^AA^ (Figure 4C). In comparison, cGAS^WT^ expression had a weak effect on CT1BA5-HSR tumors’ overall growth curves (p = 0.022); however, no individual timepoint reached significance. cGAS^AA^ had no effect on HSR tumor growth (Figure 4B). Mice with CT1BA5-HSR tumors were not protected by either cGAS^WT^ or cGAS^AA^ (Figure 4C). Therefore, ectopic cGAS expression can provide a selective therapeutic effect against ecDNA^+^ tumors.

**Figure 4.**
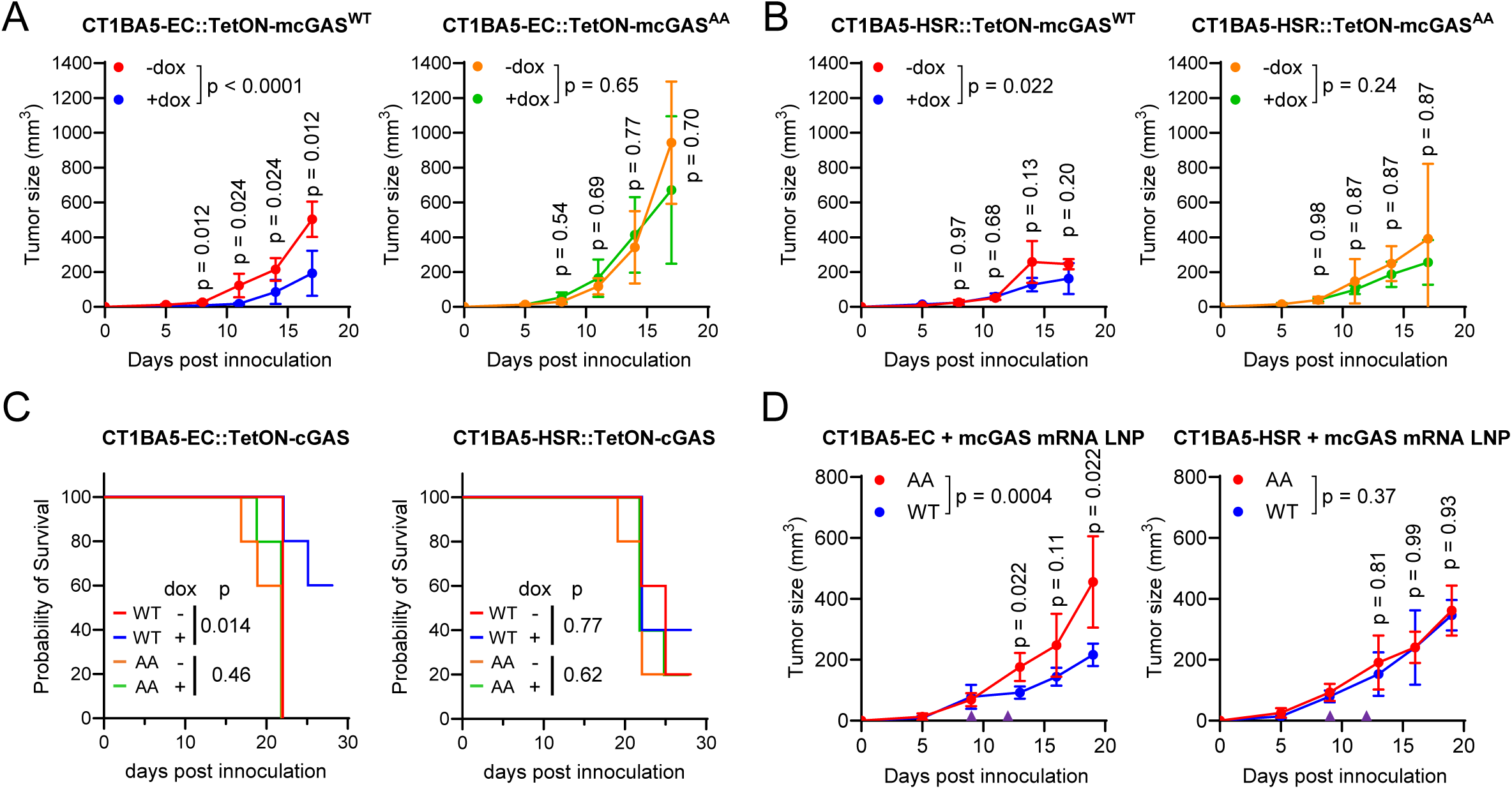
The anti-tumor effect of restoring the cGAS-STING pathway in murine ecDNA^+^ cancer cells. (**A-B**) 5x10^5^ CT1BA5-EC (**A**) or CT1BA5-HSR (**B**) cells carrying tetracycline-inducible *cGAS* (WT or catalytically dead AA) were sub-cutaneously injected into the flanks of C57BL/6J mice. The mice were fed with regular water (-dox) or water with 200 ng/mL doxycycline (+dox) and tumor growth was monitored at indicated times. (**C**) Survival curves of C57BL/6J mice that were fed with regular water (-dox) or water with 200 ng/mL doxycycline (+dox) after 5x10^5^ CT1BA5-EC or CT1BA5-HSR cells carrying tetracycline-promoter controlled cGAS (WT or catalytically dead AA) were sub-cutaneously injected into the flanks. (**D**) 2.5x10^5^ CT1BA5-EC or CT1BA5-HSR cells were sub-cutaneously allografted onto the flanks of C57BL/6J mice. LNPs carrying 5 µg mRNAs encoding WT or catalytically dead (AA) murine *cGAS* were injected intratumorally on Day 9 and Day 12 (purple triangles). Tumor growth was measured on indicated days.

We next tested the lipid nanoparticle (LNP) technology to deliver *cGAS* mRNA into the CT1BA5 tumors. *In vitro* transcribed mRNAs that contain pseudo-uridine and encode murine *cGAS* were encapsulated in LNPs and delivered to established tumors. In the CT1BA5-EC tumors, LNPs of *cGAS*^WT^ mRNA showed substantial inhibition of tumor growth compared to those treated with *cGAS*^AA^ LNPs (Figure 4D, p = 0.0004). In contrast, CT1BA5-HSR tumors treated with *cGAS*^WT^ LNPs were not different from *cGAS*^AA^ treated HSR tumors (p = 0.37). These results demonstrate the feasibility of leveraging the ecDNA to activate cGAS for cancer immunotherapy by restoring cGAS expression through mRNA LNP technology.

### The cGAS-STING pathway is a barrier to *de novo* ecDNA biogenesis

To test if cGAS-STING signaling plays a role in ecDNA biogenesis, we used the CRISPR-C^8^ technology to generate an ecDNA carrying the dihydrofolate reductase (*DHFR*) locus, which is produced after two flanking CRISPR cuts and selected with methotrexate (MTX), which kills cells without DHFR amplification^39^ (Figure 5A). This system was implemented in the human near-haploid cell line HAP1, which expresses cGAS but no detectable STING (Figure S6A). The DNA sensing capacity can be restored after ectopically expressing wild-type STING but not the R238A/Y240A mutant (Figure S6B). After MTX selection, the parental cell line gave rise to MTX-resistant colonies (Figure 5B). Similar numbers of MTX-resistant colonies were found in HAP1 cells expressing GFP or the R238A/Y240A STING mutant. In contrast, HAP1 cells expressing wild-type STING could not efficiently produce MTX-resistant colonies. We further quantified the formation of ecDNA junction and genomic scar after CRIPR-C using PCR primers targeting those specific DNA regions (Figure 5A). qPCR result showed that the formation of ecDNA and genomic scar were both strongly inhibited in HAP1 cells expressing functional STING (Figure 5C). Furthermore, the inhibition could be selectively reversed in STING-expressing HAP1 cells lacking cGAS or IRF3 (Figure 5C and Figure S6C). The result indicates that the functional cGAS-STING pathway inhibits ecDNA formation.

**Figure 5.**
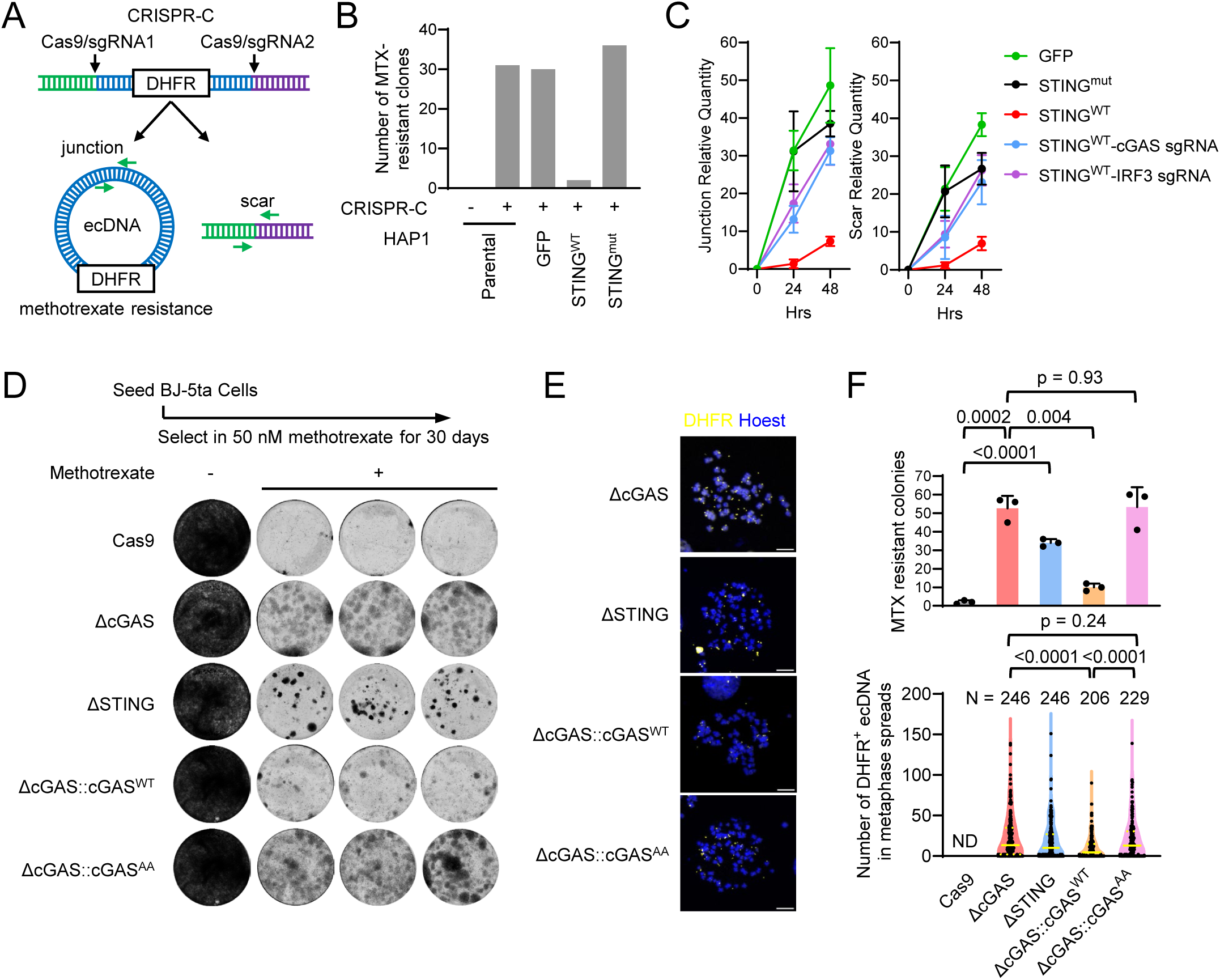
The cGAS-STING pathway restricts ecDNA formation. (**A**) Schematic of CRISPR-C to generate DHFR^+^ ecDNA. HAP1 cells were electroporated with Cas9-sgRNA RNPs targeting two sites flanking the *DHFR* locus, seeded in 6 well plate and selected in 50 nM methotrexate (+) for 4 weeks. (**B**) Bar graphs show the number of surviving HAP1 cell colonies. (**C**) The formation of ecDNA and genomic scar was quantified by qPCR at indicated time after CRISPR-C. **(D)** BJ-5ta cells were seeded in 6 well plate for one week (-) or selected in 50 nM methotrexate for 30 days (+). Live cell colonies were stained with Coomassie blue. (**E**) *DHFR*^+^ ecDNA shown by fluorescent in situ hybridization (FISH) in metaphase spreads of indicated BJ-5ta cell lines. Bars = 10 µm. **(F)** Numbers of MTX resistant colonies shown in (D) (upper) and numbers of *DHFR*^+^ ecDNA in metaphase spreads shown in (E) (bottom) for each BJ-5ta cell line as indicated.

In another approach, we selected *DHFR* ecDNA in human diploid fibroblasts BJ-5ta with MTX. Unlike cancer cells or the HAP1 cells, the cGAS-STING pathway is intact in BJ-5ta. BJ-5ta cells deficient in cGAS or STING produced much more MTX-resistant colonies than the parental WT cells that only expressed Cas9 but no specific sgRNA (Figure 5D; upper panel in 5F). Expressing wild-type cGAS in cGAS-deficient BJ-5ta cells drastically reduced the number of MTX-resistant colonies (p = 0.004), whereas the catalytic mutant cGAS^AA^ did not have a significant effect (p = 0.93, upper panel in Figure 5F). The *DHFR*^+^ ecDNA in these cells was detected by FISH (Figure 5E). cGAS^WT^ but not cGAS^AA^ reduced the number of *DHFR*^+^ ecDNA in resulting MTX-resistant cells compared to those in cGAS^KO^ cells (lower panel in Figure 5F). These results further support that cGAS-STING signaling restricts *de novo* ecDNA biogenesis.

## Discussion

The formation and propagation of ecDNA harboring oncogenes confer advantages to tumor evolution but may also represent a vulnerability for tumors if the ecDNA is sensed as foreign by the host immune system. Indeed, DNA sensing by cGAS is an important mechanism that activates the immune system, including anti-tumor immunity. In an arms race with the host immune system, many cancer cells evolve mechanisms to evade immune detection. We found that many cancer cells silence the expression of cGAS or STING and may have other mechanisms to inactivate signaling downstream of STING. Importantly, we show that restoring the expression of cGAS or STING is sufficient to reinvigorate innate immune responses in ecDNA^+^ but not ecDNA^-^ cancer cells, suggesting that cGAS is activated by ecDNA. Evidence was presented to show that cGAS binds to ecDNA in the cytoplasm and undergoes phase separation and activation. We notice that the strength of cGAS activation varied among ecDNA^+^ cell lines. This implies that there are varying levels of ecDNA in the cytosol, which are subject to the influence of total ecDNA copy numbers and the cell division process. Indeed, we show that CIP2A deficiency promotes cGAS activation, likely because CIP2A restrains ecDNA from the cytosol. Future work that aims at elucidating cellular mechanisms that tether ecDNA to mitotic chromosomes would deepen the understanding of how ecDNA normally evades cGAS-mediated sensing and enlighten strategies to target such processes to further enhance cGAS activation.

We have established an immunocompetent mouse model of ecDNA^+^ tumors to study anti-tumor immune responses *in vivo*. Using the CT1BA5 tumor model, we show that ecDNA tumors are immunologically “cold”, exhibiting poor lymphocyte infiltration and being refractory to anti-PDL1 or anti-CTLA4 immune checkpoint blockade therapies. The CT1BA5-EC tumors are also resistant to the Kras^G12D^ inhibitor MRTX1133. Interestingly, we show that these pancreatic adenocarcinoma cancer cells respond to cGAMP or cGAS expression in the tumor cells, demonstrating the efficacy of targeting the cGAS-STING pathway to rejuvenate antitumor response in these aggressive tumors. We envision that the CT1BA5 ecDNA^+^ and ecDNA^-^ tumor models will be useful research tools to further dissect how ecDNA^+^ cancer cells interact with the tumor microenvironment.

Using the CT1BA5 model, we demonstrated that activating the DNA sensing pathway in ecDNA tumors exhibits an antitumor effect. Because cGAS senses DNA in a sequence-independent manner, we propose that this presents a general strategy for targeting all ecDNA species regardless of cancer type. While the roles of the cGAS-STING pathway in tumor immunology have been extensively studied, previous investigations primarily focused on host immune cells in sensing tumor-derived DNA^40,41^. Here we provide evidence that activating this pathway selectively in tumor cells can also deliver an anti-tumor effect. Tumor-cell-derived interferons and cytokines may activate the function of dendritic cells and CD8 T cells. Tumor-derived cGAMP can also act in a paracrine manner. Furthermore, activating the cGAS-STING pathway in ecDNA^+^ cells can restrict ecDNA in a cell-autonomous manner, as we have demonstrated using the *DHFR*^+^-ecDNA-methotrexate method. Our results reveal that the cGAS-STING pathway is required and sufficient to restrict the emergence and persistence of ecDNA. Therefore, restoration of the cGAS-STING pathway in ecDNA^+^ cancer cells inhibits tumorigenesis through both tumor-cell-intrinsic and -extrinsic mechanisms.

The mRNA lipid nanoparticle technology presents a general method to ectopically express cGAS in tumors. Our data demonstrated that cGAS is selectively activated in ecDNA^+^ tumor cells. Although mRNA LNPs can only target a subset of tumor cells, the cGAS-STING pathway activates cytokines that modulate the tumor immune microenvironment. The product of cGAS, cGAMP, can propagate the signal to nearby cells via gap junction or cGAMP transporters, thus initiating a cascade of signaling events to activate an antitumor response. In parallel, we also demonstrate that decitabine, an FDA-approved anti-cancer drug, reactivates endogenous cGAS expression by reversing epigenetic silencing and subsequently elicits anti-tumor activity in ecDNA^+^ tumors. Further investigation of the mechanism of epigenetic silencing of cGAS may lead to the development of more potent and specific compounds that restore cGAS expression as anti-cancer agents.

### Limitation of the study

Several limitations should be considered when interpreting the data from this study. First, much of the mechanistic dissection of ecDNA-dependent cGAS activation was performed in a defined set of human and murine cancer cell lines. While these models span multiple tissue origins and include isogenic ecDNA–HSR pairs, they may not fully capture the heterogeneity of ecDNA organization, abundance, or intracellular dynamics present across human tumors *in vivo*. Accordingly, the extent to which cytosolic ecDNA engages cGAS across all ecDNA-positive cancers remains to be determined.

Second, although ecDNA sequences were enriched in the cytosolic fractions of ecDNA-positive cells and functionally correlated with cGAS activation, the absolute abundance of ecDNA in the cytosol is low. While cytosolic ecDNA could be visualized by FISH in a subset of cells, such events were rare, limiting quantitative assessment by imaging approaches. Consequently, our conclusions rely on the integration of biochemical fractionation, cGAMP measurements, and functional reporters. Future improvements in imaging sensitivity and resolution may enable more comprehensive visualization of cytosolic ecDNA dynamics.

Third, although promoter CpG island hypermethylation strongly correlates with cGAS silencing in multiple tumor models, regulation of the cGAS–STING pathway may also occur at additional nodes, including other pathway components. Moreover, the precise molecular mechanisms and enzymatic machinery responsible for establishing and maintaining cGAS silencing in cancer cells remain to be defined and warrant further investigation.

Finally, while our data demonstrate that an intact cGAS–STING–IRF3 pathway restricts *de novo* ecDNA biogenesis, the downstream mechanisms by which this signaling axis suppresses ecDNA formation are not yet understood. How innate immune signaling interfaces with DNA damage responses, DNA repair pathways, or chromosome segregation processes to limit ecDNA generation remains an important open question.

Despite these limitations, the convergence of genomic, biochemical, imaging, and functional *in vivo* data supports a model in which ecDNA represents a unique source of cytosolic DNA that is normally constrained by cGAS–STING signaling, and whose evasion contributes to both immune suppression and ecDNA persistence in cancer. These findings raise the exciting possibility of converting ecDNA from a selective advantage of tumors to a vulnerability that may be harnessed by the immune system to attack these aggressive tumors.

## STAR Methods

**Key resources table**

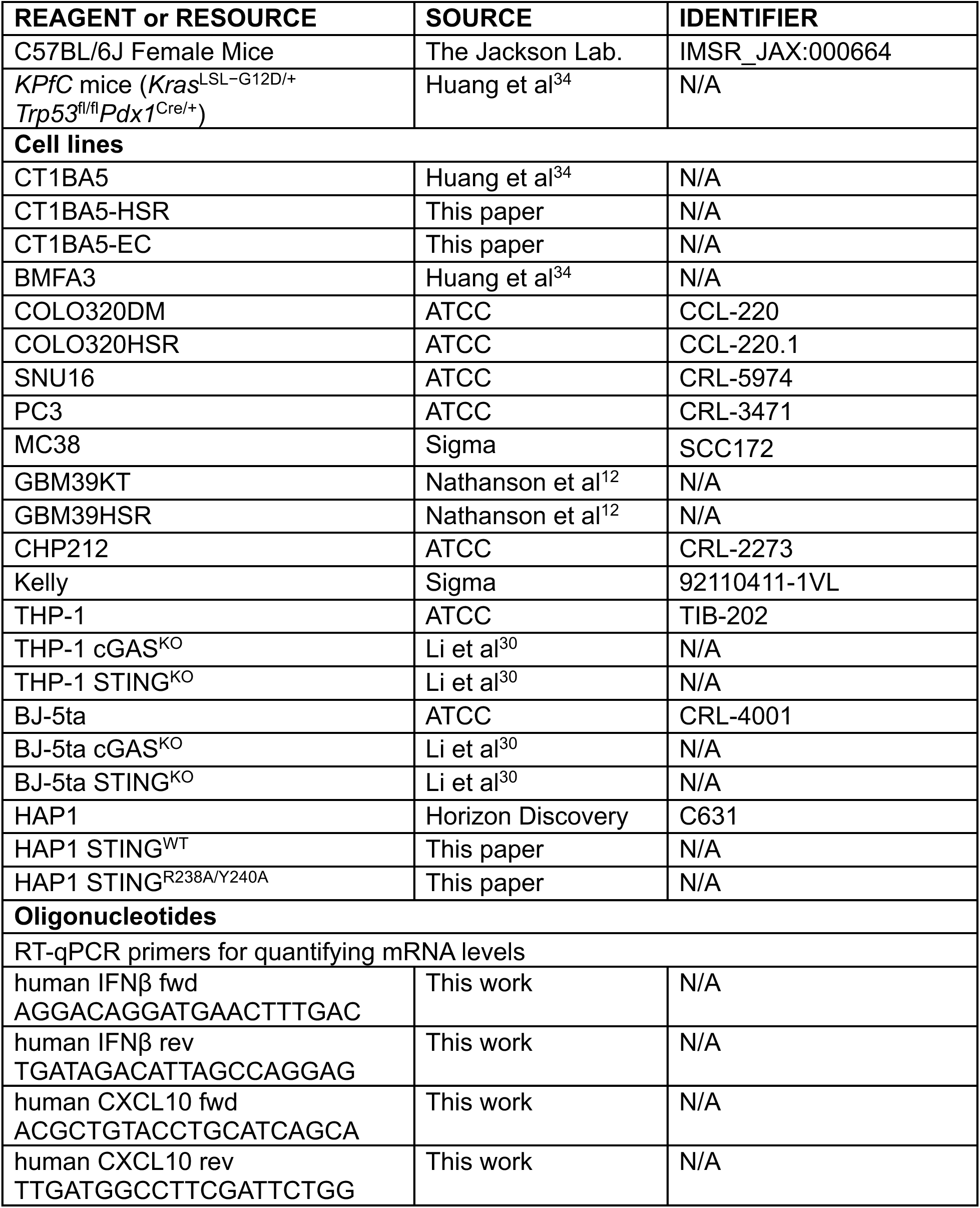
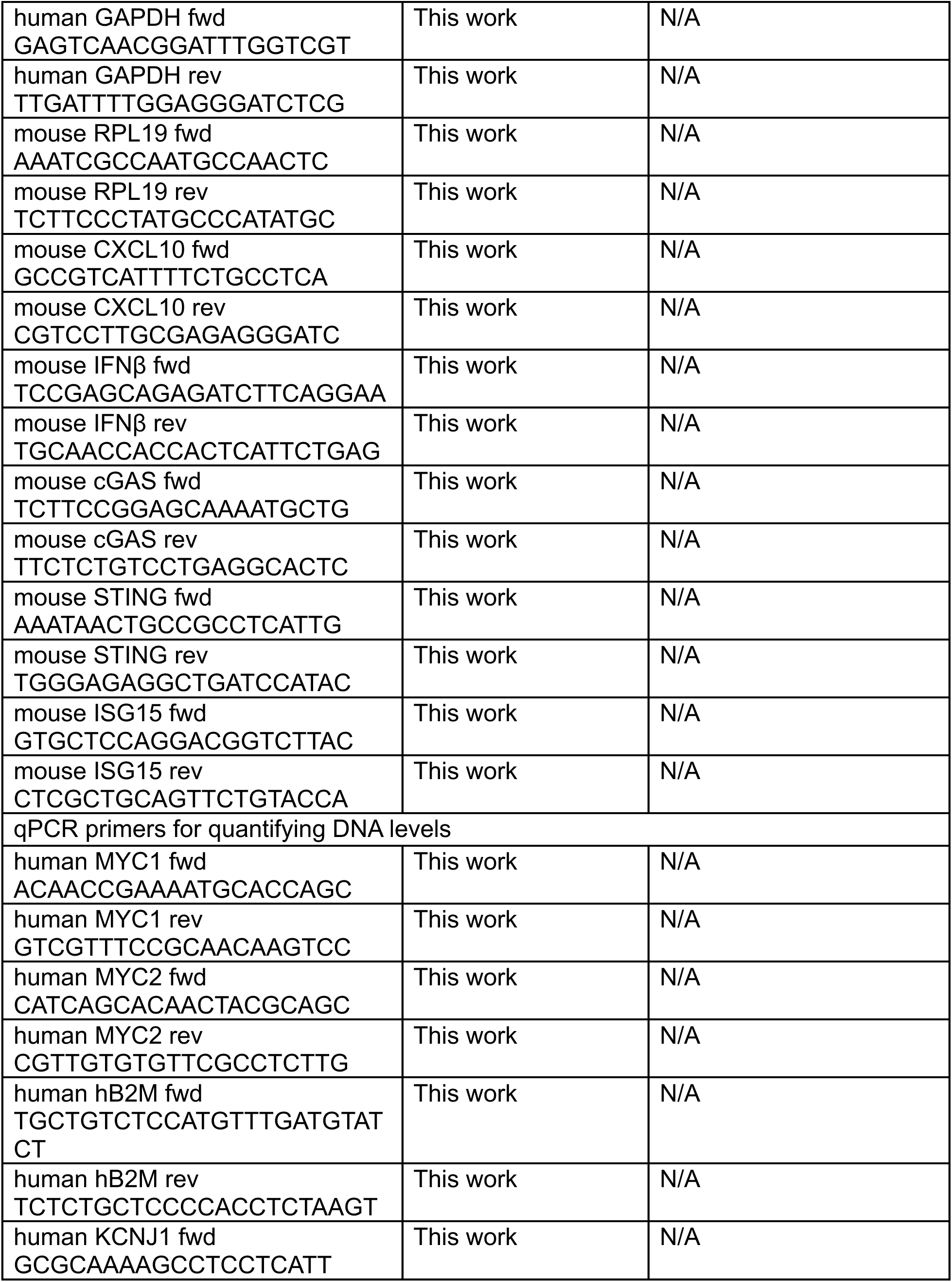
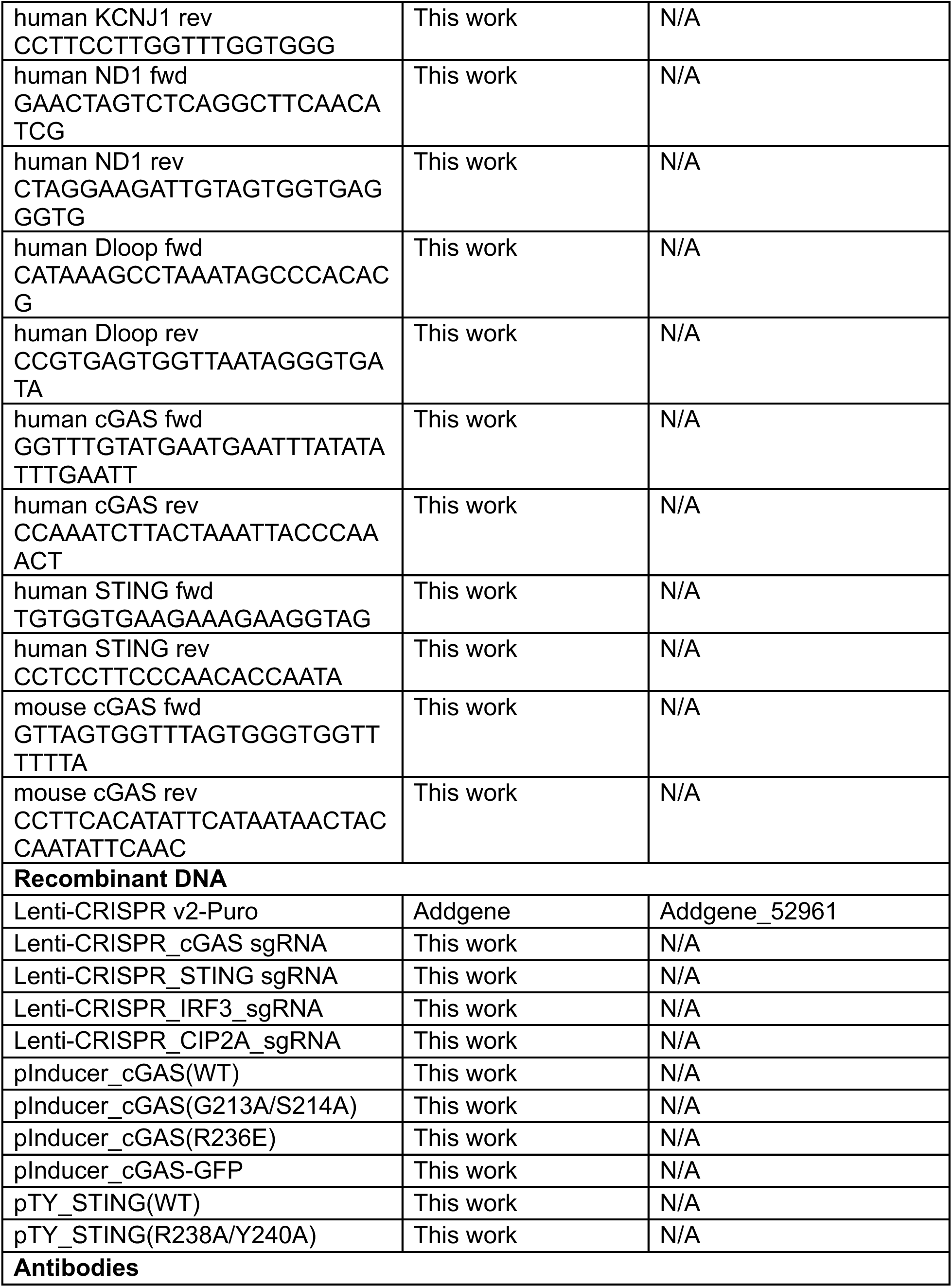
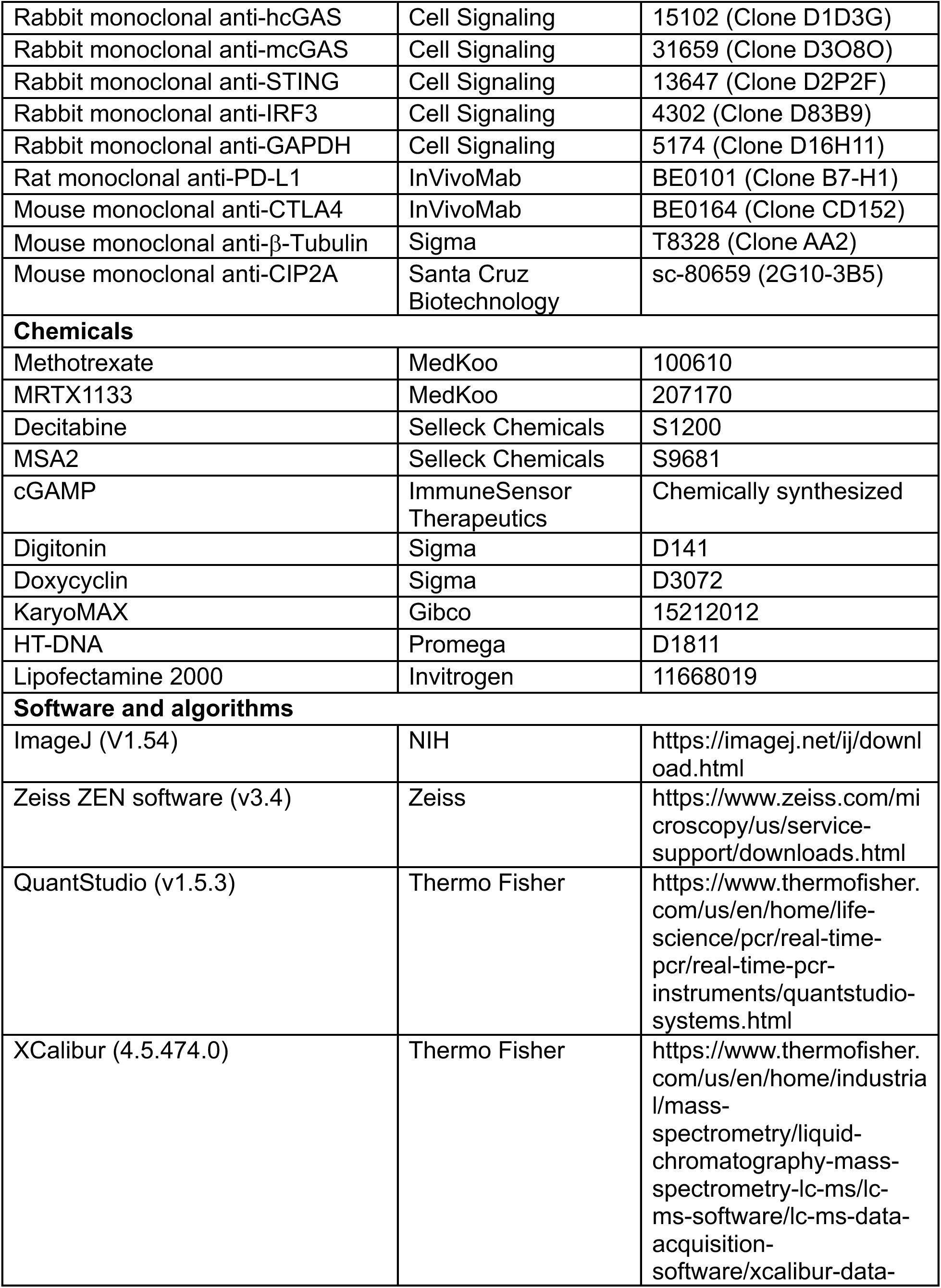
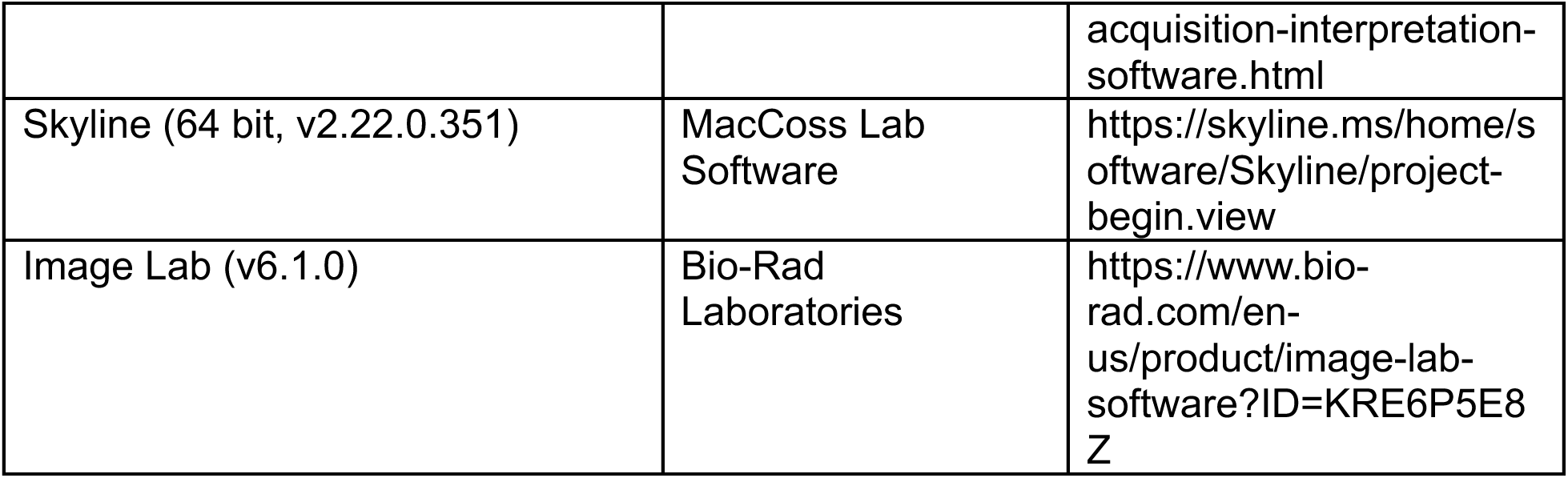

### Experimental model and study par2cipant details

#### Cell culture

Cultured cells were maintained in medium (Gibco, Thermo Scientific) supplemented with 10% fetal bovine serum (Sigma) and 1% penicillin and streptomycin in a 37°C humidified incubator with 5% CO_2_. HEK293T, BJ-5ta, CT1BA5-EC, CT1BA5-HSR, BFMA3, PC3 cells were cultured in Dulbecco’s modified Eagle’s medium (DMEM). COLO320DM and COLO320HSR cells are cultured in DMEM/Nutrient Mixture F-12. THP-1 and SNU16 cells were cultured in RPMI 1640 Medium. THP-1 Lucia cells were cultured in RPMI 1640 medium supplemented with 55 µM β-Mercaptoethanol. GBM39KT and GBM39HSR cells were cultured in DMEM/F12 supplemented with B27 supplement (Gibco), GlutaMAX, 20 ng/ml EGF, 20 ng/ml FGF, and 5 µg/ml Heparin.

#### Bisulfite sequencing

Cells were harvested by centrifugation at 400 g for 4 min. Genomic DNA was isolated using the Quick-DNA Miniprep Kit (ZYMO RESEARCH, D3025), following the manufacturer’s instructions. Subsequently, 500 ng of genomic DNA was subjected to bisulfite treatment and purification using the EZ DNA Methylation Kit (ZYMO RESEARCH, D5021). Specific regions of interest were amplified by PCR using bisulfite-specific primers (Supplementary Table 1) and Hot-Start Taq DNA Polymerase (Takara, R007A). Amplicons were purified via gel extraction with the QIAquick Gel Extraction Kit (QIAGEN, 28706) and sequenced using Premium PCR sequencing (Plasmidsaurus). DNA methylation levels were analyzed with the QUMA platform^42^. For each locus, methylation analysis was based on over 100 distinct valid reads.

#### cGAMP measurement

To measure cGAMP, 10^6^ cells were lysed in 50 µL hypotonic buffer at 95°C for 10 min. After centrifugation at 10,000 g for 10 min, 5 µL supernatant was added to 2.5×10^5^ THP-1-Lucia ISG cells seeded in a 96-well plate in 50 µL serum-free RPMI 1640 medium supplemented with 50 ng/mL recombinant perfringolysin O (PFO), and incubate at 37°C for 16 hr. 20 µL culture was mixed with 100 µL buffer 1 µM coelenterazine (Gold Biotechnology), 50 mM Tris-HCl, pH 7.0, 50 mM NaCl, 20 mM EDTA, and 20% glycerol in opaque 96-well plate. Pure cGAMP (0.3 nM to 100 nM) was used to obtain a standard curve. Measure bioluminescence with a CLARIOstar microplate reader (BMG LABTECH).

#### Reverse transcription and qPCR

Total RNA was purified from cells using the Trizol reagent (Thermo Scientific). cDNA was synthesized with the high-capacity cDNA reverse transcription kit (Applied Biosystems). The relative abundance of target transcripts was quantified by the ΔΔC_T_ method on a QuantStudio 5 Real-Time PCR instrument (Applied Biosystems) using *GPADH* (human) or *RPL19* (mouse) as the internal controls. Primers used for qPCR are listed in Supplementary Table 1.

#### Fluorescence Airyscan microscopy and FRAP assay

For live cell imaging, cells seeded in chambered glass slides (Lab-Tek) were stained with 2 µM Hoechst 33342 (Thermo Scientific) 10 min. After replacing the medium, the slides were housed in a 37°C live-cell-imaging chamber and were visualized by Airyscan microscopy on a Zeiss ASM 980 microscope equipped with Airyscan 2 and a 60× oil immersed objective. Cellular fluorescence recovery after photobleaching (FRAP) experiments were performed with a standard protocol in the ZEN software. Time-lapse images were acquired over a 14 second time course after bleaching with 0.49 second interval. Images were processed in ImageJ.

#### Subcellular fractionation and *in vitro* cGAS activity assay

Cells were lysed in a hypotonic buffer (10 mM Tris-HCl, pH 7.4, 5 mM MgCl_2_) with 1 min vigorous mixing by plastic pellet pestle in 1.5 mL tubes. Centrifuge at 1,000 g at 4°C for 10 min to obtain supernatant (S1) and pellet (P1). S1 was centrifuged at 5,000 g at 4°C for 10 min to obtain supernatant (S5) and pellet (P5). Centrifuge S5 at 10,000 g at 4°C for 10 min to obtain supernatant (S10) and pellet (P10). Resuspend pellets in hypotonic buffer of the same volume to the corresponding supernatants. The fractions were supplemented with a reaction mixture of 20 mM Tris-HCl, pH 7.4, 5 mM MgCl_2_, 0.2 mg/mL BSA, 1 mM ATP, and 1 mM ^15^N_5_-GTP (Sigma). After 1 hr incubation at 37°C, samples were boiled for 10 min and centrifuged at 20,000 g for 5 min. After spiking 80 fmol internal standard (^15^N_10_-cGAMP, in-house generated) to each sample, levels of ^15^N_5_-cGAMP were quantified by LC-MS as previously described^30^.

#### Isolate cytosolic DNA from COL320DM and COLO320HSR cells

Extract total and cytosolic DNA from COLO320DM and COLO320HSR cells to quantify the levels of nuclear, amplicon, and mitochondrial DNA in the cytosol similar to previously described^43^. For total DNA, 10^6^ cells were washed with PBS, lysed in 500 µL denaturing buffer (50 mM HEPES, pH 7.5, 150 mM NaCl, 0.1% SDS), boiled in 95°C for 10 min. For cytosolic DNA, 10^6^ cells were lysed in a digitonin buffer (50 mM HEPES, pH 7.5, 150 mM NaCl, 40 µg/mL digitonin, 1M hexylene glycol) on ice for 10 min. Supernatants were collected after centrifugation at 17,000 g for 10 min at 4°C. Total DNA and cytosolic DNA samples were both digested with Proteinase K at 55°C for 1 h, followed by phenol-chloroform isoamyl alcohol extraction. Briefly, add equal volume of phenol-chloroform isoamyl alcohol to the samples, shake vigorously for 30 seconds, centrifuge at 18,300 g for 15 min at 4°C, and transfer the supernatant to a new tube. Add 500 uL isopropanol and 1 uL glycol-blue and incubate at -20°C overnight. Centrifuge the samples at 18,300 g for 10 min at 4°C and decant the supernatant. Wash the pellet with 500 µL 70% ethanol and spin at 18,300 g for 10 min at 4°C. Decant the supernatant and air dry the pellets before dissolving them in 200 µL 10 mM Tris-HCl buffer (pH 8.0). Add 500 µg/mL RNase A and incubate at 37°C for 1 hr. Perform phenol-chloroform isoamyl alcohol extraction again and resuspend the pellet in 50 µL 10 mM Tris-HCl buffer. Quantify nuclear targets (*b2m,* kcnj1), amplicon targets (*myc1,* myc2) and mitochondrial targets (nd1 and dloop1). Primers used for qPCR are listed in Supplementary Table 1.

#### Metaphase spread and FISH

For metaphase synchronization, cells were treated with 0.1 µg/ml KaryoMAX (Gibco) for 3 h. Wash cells with PBS and suspend single cells in 75 mM KCl for 15–30 min. Add equal volume of Carnoy’s fixative (3:1 methanol:glacial acetic acid, v/v) to fix cells and wash three times with the fixative before dropping cells onto humidified glass coverslips.

Age coverslips containing fixed cells in metaphase overnight, and then equilibrate them in 2× SSC buffer, followed by 2 min dehydration in ascending ethanol series (70%, 85%, 100%). Add pre-warmed FISH probes (Empire Genomics) onto the slide and seal the coverslip with rubber cement. Denature the FISH probe and sample on a 75°C hotplate for 3 min and incubate the hybridization overnight at 37°C in a humidified chamber. On the next day, remove coverslips, wash for 2 min in 0.4× SSC at 72°C and then for 2 min in 2× SSC, 0.05% Tween-20. Stain DNA with 1 µg/ml DAPI for 2 min, wash with 2× SSC. Apply a mounting medium (VectaShield) and mount the coverslip onto a glass slide.

#### Mice and tumor allograft studies

The *KPfC* (*Kras^LSL-G12D/+^*; *Trp53^fl/fl^*; *Pdx1^Cre/+^*) mice were generated as previously described.^22^ The *KPfC* mouse had a pure C57BL/6 genetic background. All other mice used in this study were purchased from The Jackson Laboratory (6–9-week-old female C57BL/6J, strain #000664). All mice used in this study were maintained under specific pathogen-free conditions in the animal facility of the University of Texas Southwestern Medical Center at Dallas according to experimental protocols approved by the Institutional Animal Care and Use Committee. Cancer cells were subcutaneously injected into the flank of mice. Tumor treatments include: 200 µg anti-PD-L1 antibody (InVivoMab), 100 μg anti-CTL4 antibody (InVivoMab), 30 mg/kg MRTX1133 (MedKoo, 207170), 10 μg cGAMP, 50 ng recombinant murine IL12 p70 (PeproTech), and 20 mg/kg decitabine (Selleck Chemicals, S1200). Tumor sizes were measured every 2 or 3 days and calculated by (length × width × height × π)/6.

#### mRNA lipid nanoparticles

mRNA encoding murine *cGAS* was generated by *in vitro* transcription. A pcDNA3.1 plasmid containing the coding sequence of murine *cGAS* downstream the T7 promoter was linearized by NotI digestion and purified by phenol-chloroform precipitation. mRNA was synthesized from 50 µg/mL linearized plasmid with 8 U/µL T7 RNA Polymerase (NEB) in 40 mM Tris-HCl (pH 8.0), 16.5 mM magnesium acetate, 2 mM spermidine, 10 mM DTT, 0.002% Triton X-100, 5 mM ATP, 5 mM GTP, 5 mM CTP, 5 mM N1-methylpseudouridine-5’-triphosphate, 4 mM m⁷G(5′)ppp(5′)(2′-O-methyladenosine)pG as the 5′ cap (CleanCap AG 3’OMe, TriLink Biotechnologies), 1 U/µL murine RNase inhibitor (NEB) and 0.002 U/µL inorganic pyrophosphatase (NEB) at 37°C for 3 hrs. The synthesized mRNA was purified using phenol-chloroform precipitation and resuspended in nuclease-free water. The integrity and size of the RNA were verified by agarose gel electrophoresis and concentration measured by a NanoDrop spectrophotometer.

We used the NanoAssemblr Ignite microfluidic mixing device (Precision NanoSystems) to manufacture mRNA-Lipid Nanoparticles (LNPs) at a total flow rate of 12 mL/min and a flow rate ratio of 3:1 between the aqueous and the ethanol phases. The aqueous phase contained mRNA (0.8 mg/mL) in 25 mM sodium acetate buffer (pH 4.0). The ethanol phase contained the lipid mixture of ionizable cationic lipid SM-102, 1,2-Distearoyl-sn-glycero-3-phosphocholine (DSPC), cholesterol, and PEG-lipid DMG-PEG2000 at a molar ratio of 50:10:38.5:1.5, respectively. All lipids were purchased from MedKoo Biosciences. The resulting LNPs were diluted in and dialyzed against PBS (pH 7.4) to remove ethanol. As quality control of LNPs, we assessed their particle size and polydispersity index (PDI) by dynamic light scattering using a Zetasizer Nano ZS (Malvern). The RiboGreen RNA assay (Thermo Fisher) with and without Triton X-100 was used to measure encapsulation efficiency. LNPs were stored at 4°C and used within 2 weeks for functional studies.

#### Statistics

All experiments were performed three times. Statistical analyses are detailed in the corresponding figure legends. For data that were normally distributed and exhibited homoscedasticity, Student’s t-test or one-way ANOVA was used. Otherwise, the corresponding statistical analysis was indicated in the figure legend. All tests were two-sided. Violin plots display the median and interquartile range (IQR). Data points and error bars shown on bar plots represent the mean ± standard error of the mean (SEM).

## Data availability

All sequencing data will be uploaded to the NIH Sequence Read Archive repository and made publicly accessible upon the acceptance of this manuscript.

## Author contributions

Tuo Li: Conceptualization, Methodology, Investigation, Validation, Formal analysis, Visualization, Writing - Original Draft. Qing-Lin Yang: Investigation, Validation, Formal analysis, Visualization, Writing - Review & Editing. Kailiang Qiao: Investigation, Writing - Review & Editing. Anli Zhang: Investigation. Chenglong Sun: Investigation. Huocong Huang: Investigation. Paul Mischel: Investigation, Resources, Writing - Review & Editing, Funding acquisition. Sihan Wu: Conceptualization, Methodology, Resources, Formal analysis, Writing - Review& Editing, Funding acquisition, Supervision. Zhijian Chen: Conceptualization, Methodology, Resources, Formal analysis, Writing - Review& Editing, Funding acquisition, Visualization, Supervision.

## Acknowledgement

We thank Matthew Jones (Stanford), Jens Luebeck, Kyra Fetter, Vineet Bafna (UCSD) and members of the Chen Laboratory for discussions. We acknowledge the assistance of Yan Liu and Ishmael Dehghan in mass spectrometry. Work in Chen laboratory is supported by grants from the National Cancer Institute (R01CA299257) and the Welch Foundation (I-1389). Work in Wu laboratory is supported by the Cancer Prevention and Research Institute of Texas (CPRIT, RR210034) and the American Cancer Society (CAT-24-1379043-01-CAT). Huocong Huang is supported by the National Cancer Institute (R00 CA252009). This work was delivered as part of the eDyNAmiC team supported by the Cancer Grand Challenges partnership funded by Cancer Research UK (P.S.M CGCATF-2021/100012, Z.J.C., S.W. CGCATF-2021/100023) and the National Cancer Institute (P.S.M. OT2CA278688, Z.J.C., S.W. OT2CA278683). Zhijian J. Chen is an investigator of the Howard Hughes Medical Institute.

---

**Figure S1.**
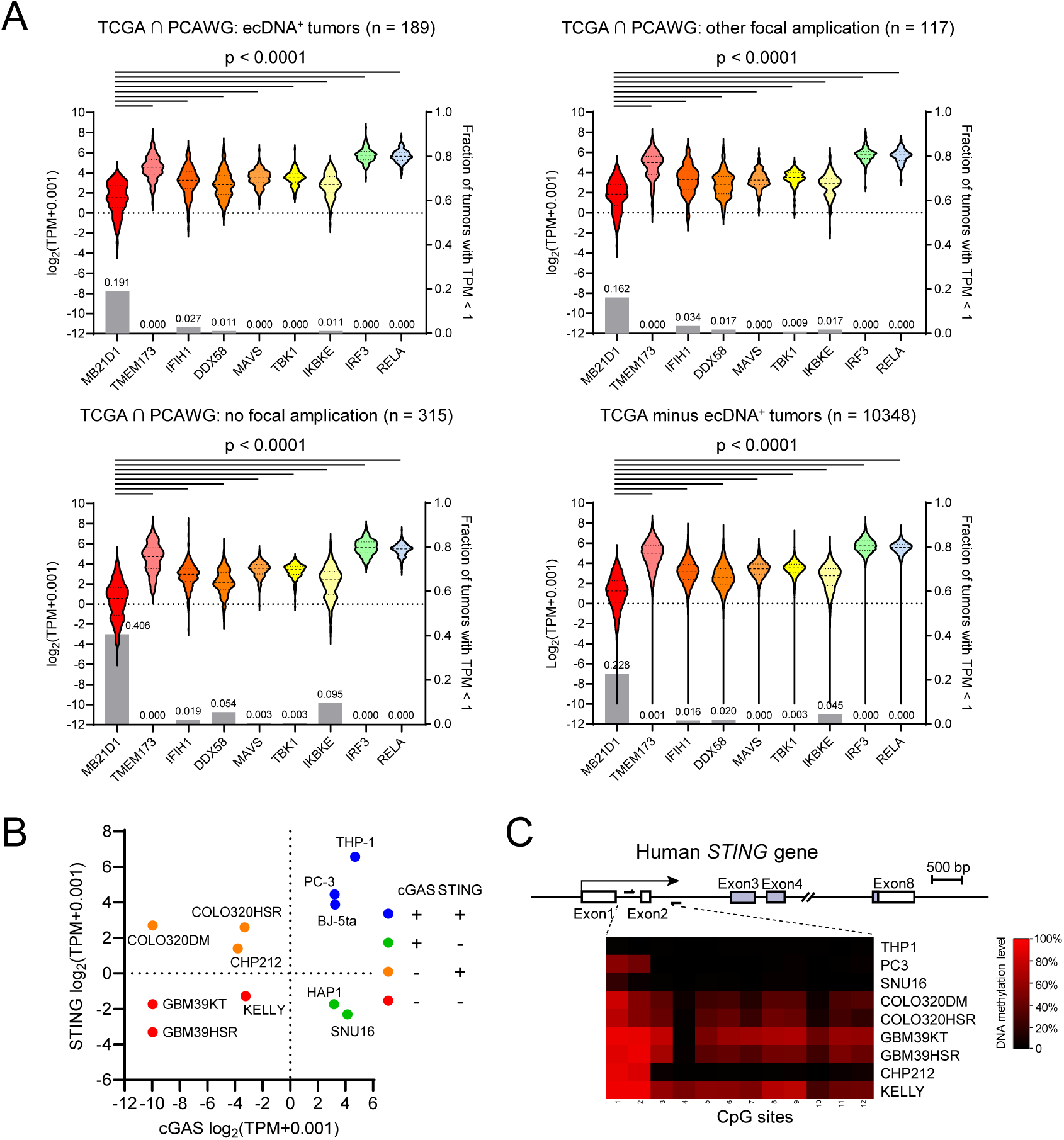
The cGAS-STING pathway is frequently defective in ecDNA^+^ cancer cells. (**A**) mRNA levels of key components of nucleic acid sensing pathways in overlapping TCGA and PCAWG tumor samples with ecDNA (*upper-left*), with other focal amplifications—BFB, non-cyclic complex or linear amplification but without ecDNA (*upper-right*), and without detected focal amplifications (*bottom-left*). (*bottom-right*) All TCGA PANCAN samples subtract ecDNA^+^ samples. (**B**) *cGAS* and *STING* mRNA levels in selected human cell lines. (**C**) The schematics show the location of the CpG rich region of human *STING* gene. The heatmaps show the DNA methylation levels of individual CpG sites as measured by bisulfite sequencing.

**Figure S2.**
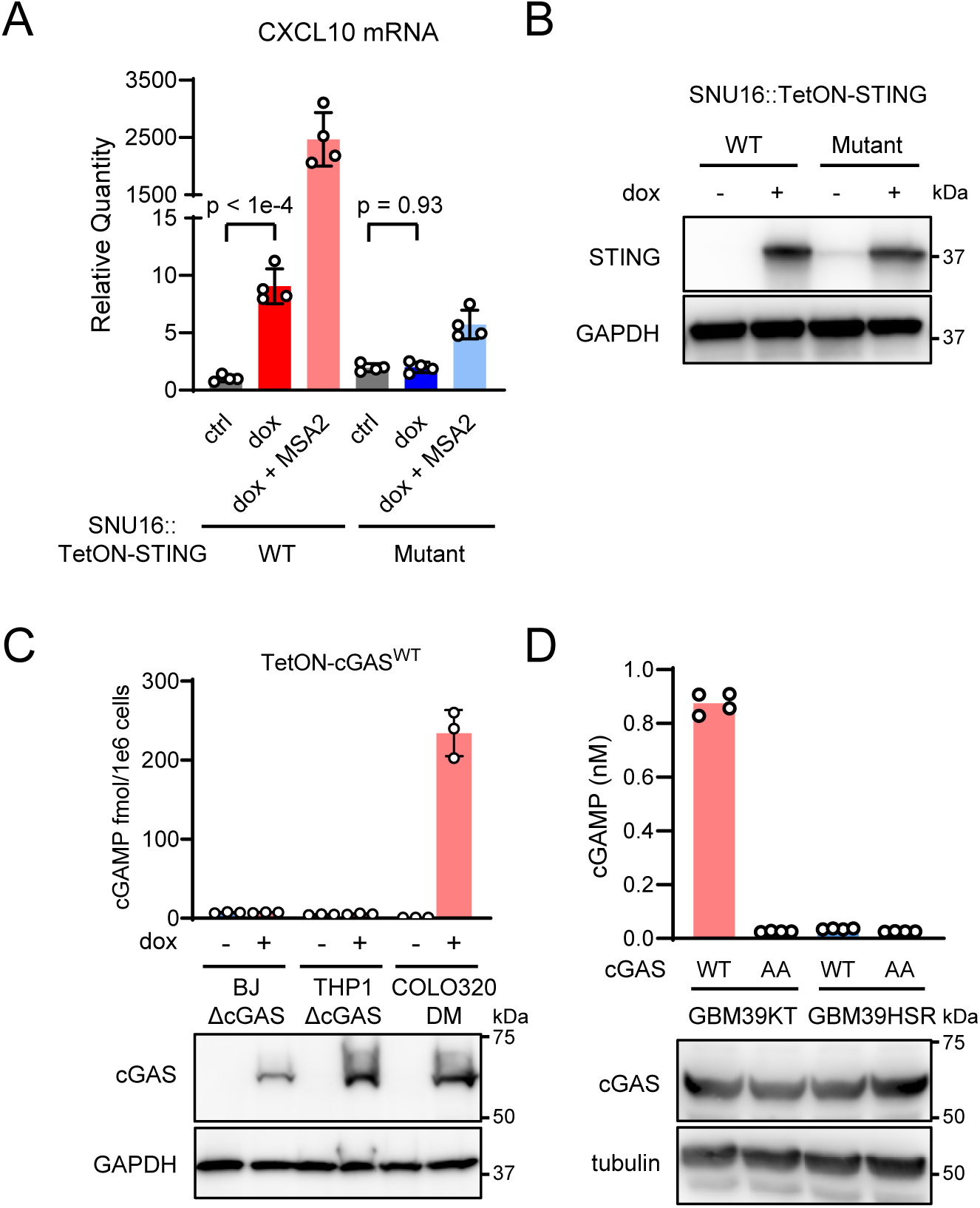
Immune activation by restoring the cGAS-STING pathway in human ecDNA^+^ cancer cells. (**A&B**) SNU16 cells (ecDNA^+^) were stably transduced with tetracycline-inducible *STING* constructs: wild-type (WT), the cGAMP-binding deficient mutant R238A/Y240A. After 0.1 µg/mL doxycycline induction for 24 hrs, *CXCL10* mRNA levels were measured by RT-qPCR (**A**) and STING protein levels determined by immunoblotting (**B**). (**C**) cGAMP levels in human BJ-5ta, THP-1, and COLO320DM cells induced to express cGAS^WT^ with doxycycline. (**D**) cGAMP levels in human glioblastoma cell lines GBM39KT (ecDNA^+^) and GBM39HSR (ecDNA^-^) expressing cGAS^WT^ or the inactive mutant cGAS^AA^ (G213A/S214A).

**Figure S3.**
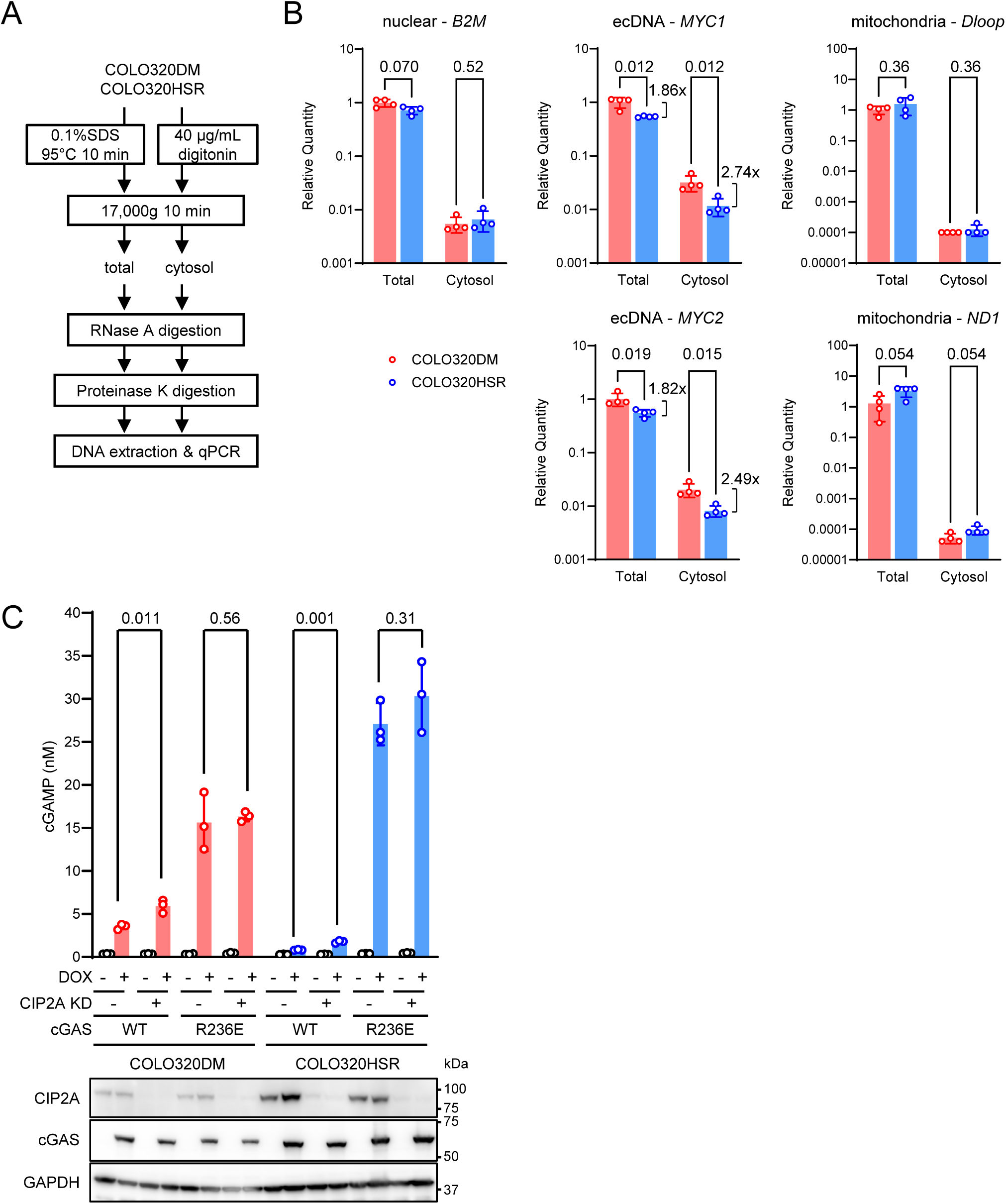
cGAS is activated by cytoplasmic ecDNA. (**A**) The workflow for isolating DNA from total or cytosolic lysate of COLO320DM and COLO320HSR cells, and for quantification of selected DNA targets. (**B**) Relative quantities of selected DNA sequences representing the nuclear genome (*B2M*), ecDNA (*MYC1* and *MYC2*), or mitochondria genome (*Dloop* and *ND1*) in fractions obtained in (A). Data are normalized to the total levels of nuclear target *KCNJ1*. (**C**) CIP2A was knocked out by Cas9-sgRNA in COLO320DM and COLO320HSR cells. cGAMP levels were quantified after inducing cGAS expression with doxycycline.

**Figure S4.**
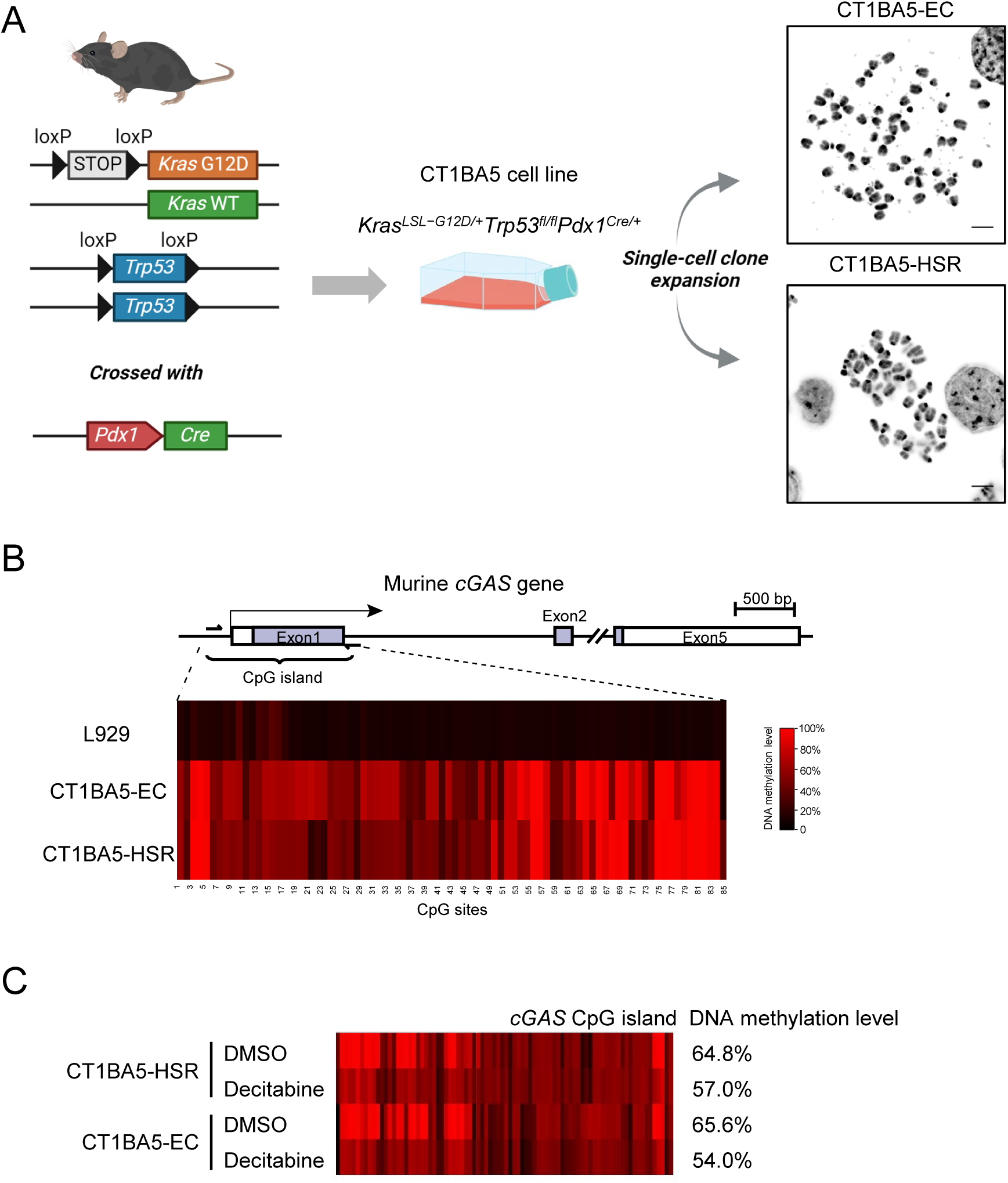
Isolation of an ecDNA^+^ cell line from the *KPfC* mice with pancreatic adenocarcinoma. **(A)** The KPfC *Kras^LSL−G12D/+^Trp53^fl/fl^Pdx1^Cre/+^*GEMM on the C57BL/6 background was generated by intercrossing *Kras^LSL−G12D/+^Trp53^fl/fl^*mice with Pdx1-Cre transgenic mice. The dual alteration of *Kras* and *Trp53* drives spontaneous initiation and progression of pancreatic ductal adenocarcinoma (PDAC). A primary pancreatic cancer cell line CT1BA5 was isolated from late-stage tumor and subsequently subcloned to generate two clonal cell lines: CT1BA5-EC (ecDNA^+^) and CT1BA5-HSR (focal amplification of *Kras* as HSR) (co-submitted manuscript by Qiao et al^35^). FISH in metaphase spread show focal amplification of *Kras*. (**B**) The upper schematic shows the location of the CpG island in the Exon 1 of murine *cGAS* gene. The bottom heatmap shows the DNA methylation levels of individual CpG sites as measured by bisulfite sequencing. (**C**) Bisulfite sequencing of *cGAS* CpG island in CT1BA5-EC and HSR cells after 10 μM decitabine or DMSO treatment for 48 hrs.

**Figure S5.**
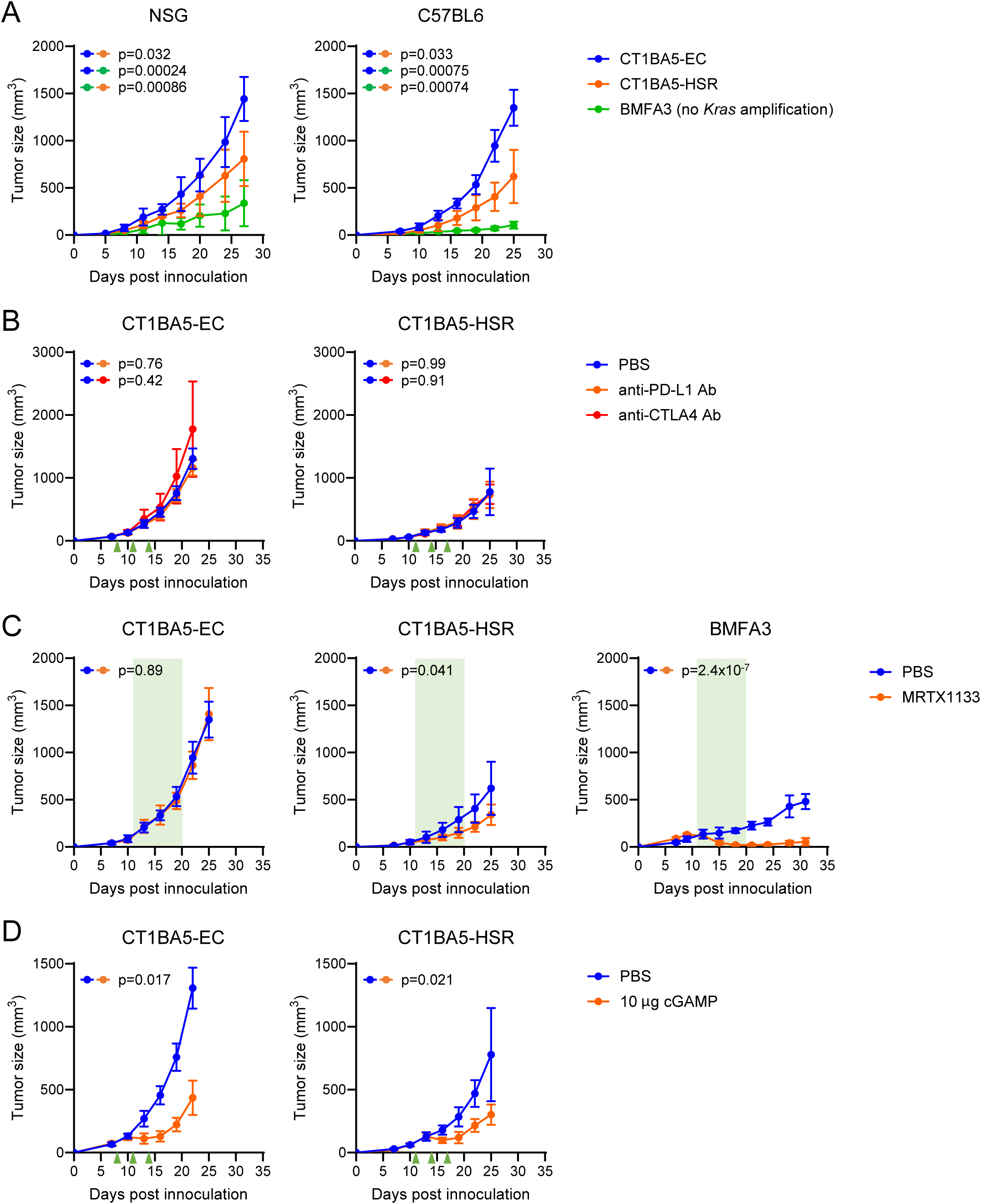
The CT1BA5 tumors are resistant to targeted therapy and immune checkpoint blockade but are sensitive to pharmacological activation of the cytosolic DNA sensing pathway. (**A**) Growth curves of CT1BA5-EC, CT1BA5-HSR and BMFA3 subcutaneous tumors in C57BL/6 (n=7-10) and NSG mice (n=6). (**B**) Growth curves of CT1BA5-EC and CT1BA5-HSR subcutaneous tumors after intraperitoneal treatment with anti-PD-L1 (200 μg) or anti-CTLA4 (100 μg) antibody at indicated times (n=4-6) (**C**) Growth curves of CT1BA5-EC, CT1BA5-HSR and BMFA3 subcutaneous tumors after intratumoral injections of MRTX1133 (30 mg/kg) twice a day. (**D**) Growth curves of CT1BA5-EC and CT1BA5-HSR subcutaneous tumors after intratumoral injections of cGAMP (10 μg) (n=4-6).

**Figure S6.**
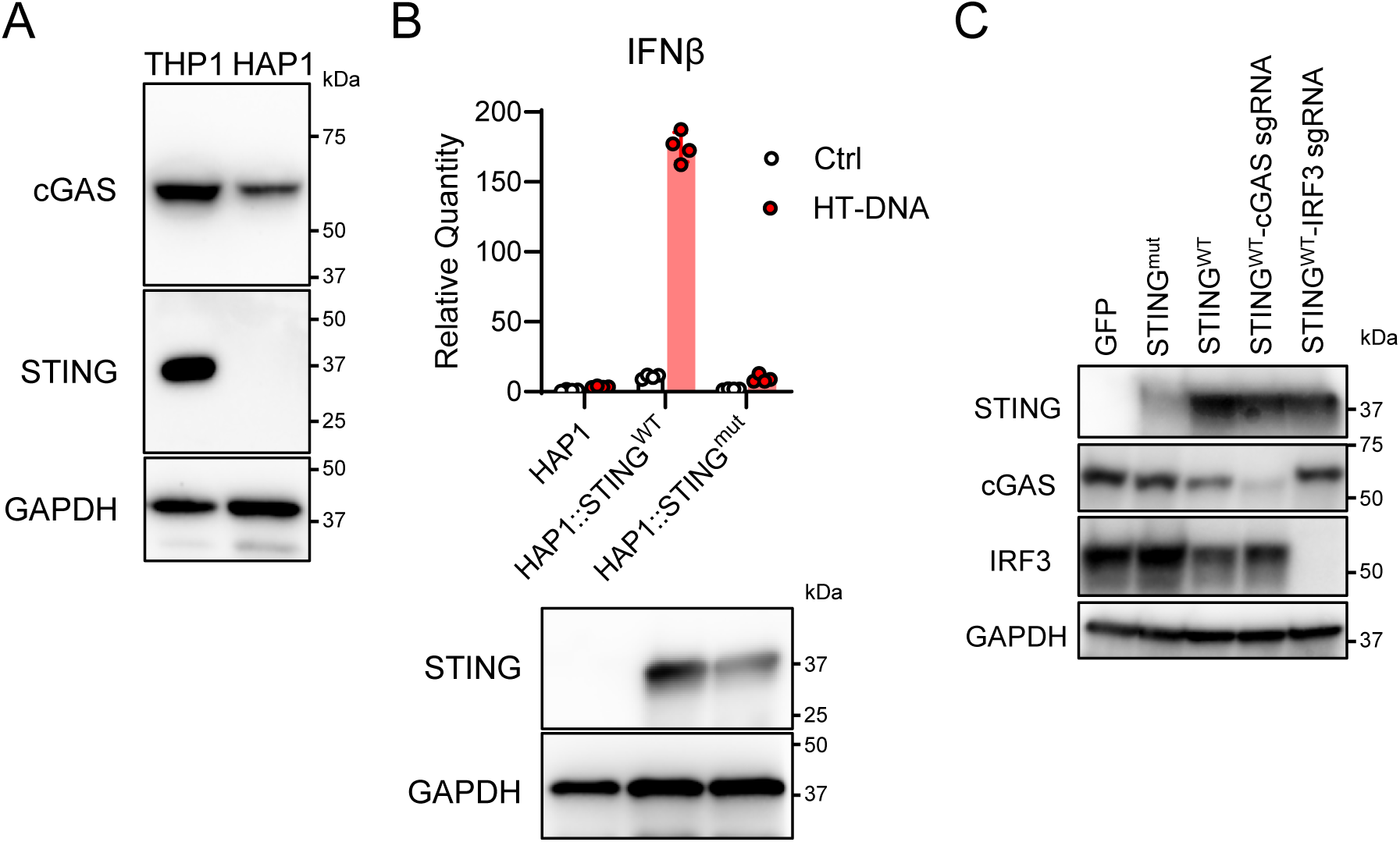
Supporting data for the roles of cGAS-STING pathway in restricting ecDNA formation. (**A**) Immunoblotts show protein levels of cGAS and STING in human HAP1 cells. (**B**) HAP1 cell lines were stably transduced with wild-type *STING* (WT) or the cGAMP-binding-deficient R238A/Y240A mutant (mut). Cells were transfected with HT-DNA for 6 hrs and the levels of *IFNβ* transcript were quantified by RT-qPCR. (**C**) Immunoblots show levels of cGAS and IRF3 after CRISPR-mediated knockdown in HAP1 cells.

